# Photoaffinity ligand of Cystic Fibrosis corrector VX-445 identifies SCCPDH as an off-target

**DOI:** 10.1101/2025.02.22.639665

**Authors:** Minsoo Kim, Kwangho Kim, Jesun Lee, Lea A. Barny, LaToya Scaggs, Ian Romaine, KyuOk Jeon, Simona G. Codreanu, Stacy D. Sherrod, John A. McLean, Anjaparavanda P. Naren, Gary A. Sulikowski, Lars Plate

**Affiliations:** Department of Chemistry, Vanderbilt University, Nashville, TN, United States; Program in Chemical and Physical Biology, Vanderbilt University, Nashville, TN, United States; Vanderbilt Institute of Chemical Biology, Vanderbilt University, Nashville, Tennessee 37232, United States; Division of Pulmonary Medicine and Critical Care, Cedars-Sinai Medical Center, Los Angeles, CA, United States; Center for Innovative Technology, Vanderbilt University, Nashville, TN, United States; Department of Biological Sciences, Vanderbilt University, Nashville, TN, United States; Department of Pathology, Microbiology and Immunology, Vanderbilt University Medical Center, Nashville, TN, United States

**Keywords:** cystic fibrosis (CF), cystic fibrosis transmembrane conductance regulator (CFTR), Elexacaftor, VX-445, VU439, photoaffinity ligand (PAL), mass spectrometry (MS), target identification, saccharopine dehydrogenase-like oxidoreductase (SCCPDH), metabolomics

## Abstract

Cystic fibrosis (CF) pharmacological correctors improve Cystic Fibrosis transmembrane conductance regulator (CFTR) protein trafficking and function. Several side-effects from these correctors and adverse drug interactions have been reported, emphasizing the need to understand off-targets. We synthesized VU439, a functionalized photoaffinity ligand (PAL) of VX-445. Chemoproteomics analysis by mass spectrometry (MS) was used to identify crosslinked proteins in CF bronchial epithelial cells expressing F508del CFTR. We identified saccharopine dehydrogenase-like oxidoreductase (SCCPDH), an uncharacterized putative oxidoreductase, as a VX-445 specific off-target. We then characterized changes in the metabolomic profiles of cells overexpressing SCCPDH to determine the consequence of VX-445 binding to SCCPDH. These data show dysregulation of amino acid metabolism and a potential inhibitory activity of VX-445 on SCCPDH. The identified off-target may explain exacerbation of psychological symptoms observed in the clinic, thus emphasizing the need for further optimization of correctors.

## Introduction

Cystic fibrosis (CF) is a prevalent genetic disorder caused by mutations in the cystic fibrosis transmembrane conductance regulator (CFTR) protein^1^, a cyclic adenosine monophosphate (cAMP)-dependent anion channel that conducts chloride and bicarbonate across epithelial apical membranes of multiple exocrine organs. Many people with CF benefit from combination therapies developed over the last decade. Pharmaceutical correctors bind and stabilize CFTR for proper maturation and function at the cell surface^3,4^. Among available correctors, the second-generation corrector VX-445 (Elexacaftor) can treat a wide range of CF-causing variants including the most common F508del variant^4–6^. Several side-effects from these correctors such as hepatotoxicity, abdominal pain, severe rashes, depression, etc., and importantly, adverse drug interactions leading to liver damage have been reported, emphasizing the need to understand the etiology of these effects that potentially arise from off-targets^7–11^.

Off-target activity of pharmaceutical drugs that result from incomplete understanding of drug targets may cause undesirable adverse effects to patients or in severe cases, discontinuation of the drug, burdening patients with unforeseen difficulties^12^. For example, painkillers, anti-inflammatory modulators, and weight loss medications were withdrawn from market due to severe side effects unaccounted during approval^13,14^. Identifying the complete range of drug targets will aid comprehension of potential adverse effects, alternative mechanisms of action, and drug repurposing (**Figure 1A**).

**Figure 1.**
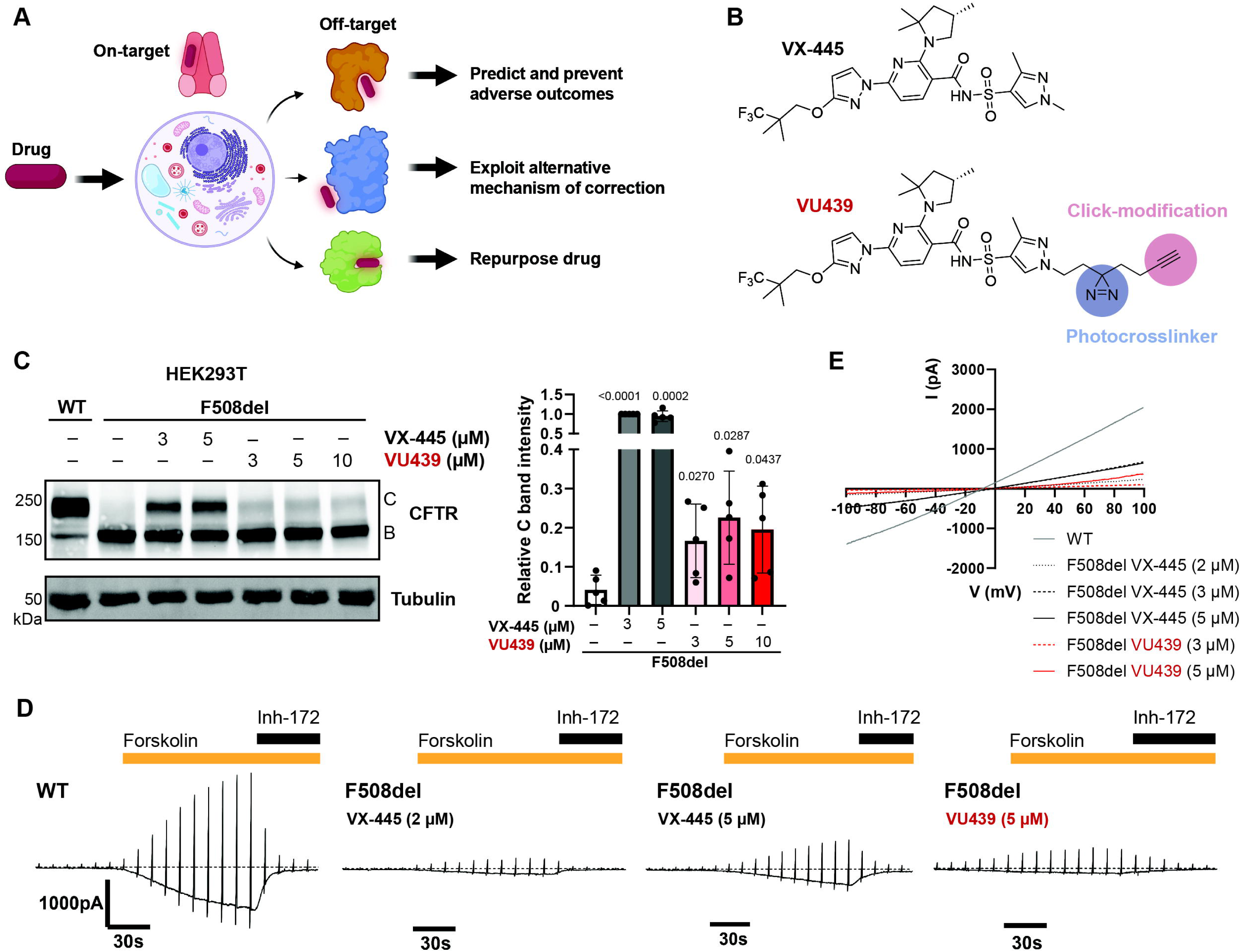
Photoaffinity ligand retains rescue of F508del CFTR trafficking and function. **A.** Understanding the complete drug target spectrum allows prediction and prevention of adverse effects and repurposing of drug towards off-targets. **B.** Molecular structures of VX-445 and photoaffinity ligand VU439. VU439 contains an alkyl diazirine moiety for UV induced photocrosslinking and a terminal alkyne for click-modifications. **C.** Representative immunoblot image showing F508del CFTR correction by VX-445 and VU439 in HEK293T cells transiently transfected with CFTR plasmids. Cells were treated with respective compound concentrations or vehicle (DMSO) for 24 h before collection. Quantification of CFTR C bands shown as mean ± SD (n = 5). VU439 shows retained correction of fully glycosylated F508del CFTR. Quantified band C values in all conditions were normalized to F508del treated with VX-445 at 3 μM. Tubulin is shown as loading control. Statistical differences were computed via one-way ANOVA with Geisser-Greenhouse correction and post-hoc Dunnett’s multiple comparisons testing against F508del treated with vehicle. P-values as shown. **D.** Representative electrophysiology traces measured by whole cell patch clamp on HEK293 cells transiently expressing WT or F508del CFTR. Data showing whole cell current (pA) during stimulation with forskolin (20 μM) and inhibition by CFTR inh-172 (20 μM). VX-445 (2 µM and 5 µM) restored F508del CFTR ion current when stimulated with forskolin. VU439 (5 µM) similarly restored ion current, showing retained functional correction. **E.** Representative I-V curve from D.

CF drugs were functionalized by other groups to confirm binding to CFTR. Sinha and co-workers functionalized VX-809 (Lumacaftor) with a terminal alkyne and showed enrichment of non-covalently bound CFTR through click chemistry and pulldowns^15^. Furthermore, Laselva and co-workers utilized a trifluoromethyl diazirine photoactivable analog of VX-770 (Ivacaftor) to determine the binding site of VX-770 on CFTR using mass spectrometry^16,17^. Thus, we sought to leverage the profiling capabilities of photoaffinity probes to characterize binding to protein targets.

Photoaffinity ligands (PAL) contain a light-activated moiety allowing covalent crosslinking during binding interactions with proteins. This tool, coupled with chemical proteomics, has been applied to identify binding of small molecules on target proteins in various biological contexts^18–21^.

Herein a functionalized photoaffinity ligand of VX-445 was synthesized that harbors a minimalist alkyl diazirine for covalent photo-crosslinking and a terminal alkyne handle for various click reaction mediated modifications (VU439). We demonstrate VU439 retains its activity as a CFTR corrector. Once the probe was crosslinked to its bound proteins in CF Bronchial Epithelial cells *in situ,* the crosslinked targets were enriched via affinity purification and characterized using LC-MS/MS to identify specific off-targets. These data identified saccharopine dehydrogenase-like oxidoreductase (SCCPDH) as an off-target of VX-445. Metabolomics data exposed two canonical pathways perturbed by overexpression of SCCPDH and effect size was further intensified by the addition of VX-445, showing its counteractivity against SCCPDH-induced changes. These findings highlight the utility of PALs in the identification of off-targets of clinically relevant compounds, as exemplified here by a widely used CF drug.

## Results

### I. Photoaffinity ligand retains rescue of F508del CFTR trafficking and function

We synthesized VU439, a functionalized photoaffinity ligand (PAL) analog of VX-445 (**Figure 1B, Figure S1A**). We developed a convergent synthesis route, opting to modify the N-methyl group of the pyrazole to contain an alkyl diazirine and a terminal alkyne. We expected minimal perturbations in binding based on the VX-445 bound cryo-EM structure^2^ where the pyrazole ring is partly solvent exposed and can likely rotate (**Figure S1B**).

We first assessed the activity of VU439 by treating cells and measuring mature CFTR rescue by immunoblotting as indicated by a complexly glycosylated post-Golgi CFTR (C band) at ∼170 kDa. In HEK293T cells transiently expressing F508del CFTR, a trafficking and function deficient variant, VU439 showed a small but significant increase in trafficking as measured by the intensity of C band (**Figure 1C**). Notably, correction of F508del CFTR trafficking was saturated at the 3 µM dose of either VX-445 or VU439.

To determine rescue of channel function, we then measured CFTR channel activity upon treatment with VU439. We performed whole cell patch clamp assays to measure ion conductivity in HEK293 cells expressing wild-type (WT) or F508del CFTR (**Figure 1D and 1E**). Briefly, cells were exposed to forskolin to induce channel opening and then inhibited with CFTR selective inhibitor-172 while measuring whole cell current to determine the activity specific to CFTR. Consistent with trafficking improvement, treatment of F508del CFTR with VU439 led to a dose-dependent increase in ion conductivity upon forskolin stimulation. VU439 at a 5 µM dose showed a comparable improvement of F508del CFTR function to VX-445 at a 2 µM dose. Even though the VU439 efficacy and potency were reduced compared to the parent compound, these data suggest VU439 retains activity for binding to mutant CFTR to rescue trafficking and function. Therefore, VU439 is suitable for off-target identification. We next sought to use VU439 to identify protein binding partners.

### I. SCCPDH was identified as an off-target of VX-445 via affinity purification mass spectrometry

To identify off-targets of CFTR corrector VX-445, we treated TetON inducible Cystic Fibrosis Bronchial Epithelial (CFBE) cells expressing F508del CFTR with VU439 (**Figure 2A**). To evaluate specificity, we included a competition condition where ten-fold access VX-445 was added to cells prior to VU439 treatment. Cells were irradiated with UV light to induce diazirine-mediated crosslinks. Lysates were obtained to further derivatize VU439 using click chemistry with a trifunctional probe containing an azide click receptor, a rhodamine fluorophore (TMR) for visualization, and desthiobiotin for affinity capture. We then enriched desthiobiotin-TMR modified proteins using streptavidin bead and resolved by SDS-PAGE to confirm VU439 dependent labeling of off-target proteins (**Figure 2B**). By SDS-PAGE, we observed no apparent disappearance of bands upon addition of excess VX-445, indicating the need for detection by quantitative mass spectrometry.

**Figure 2.**
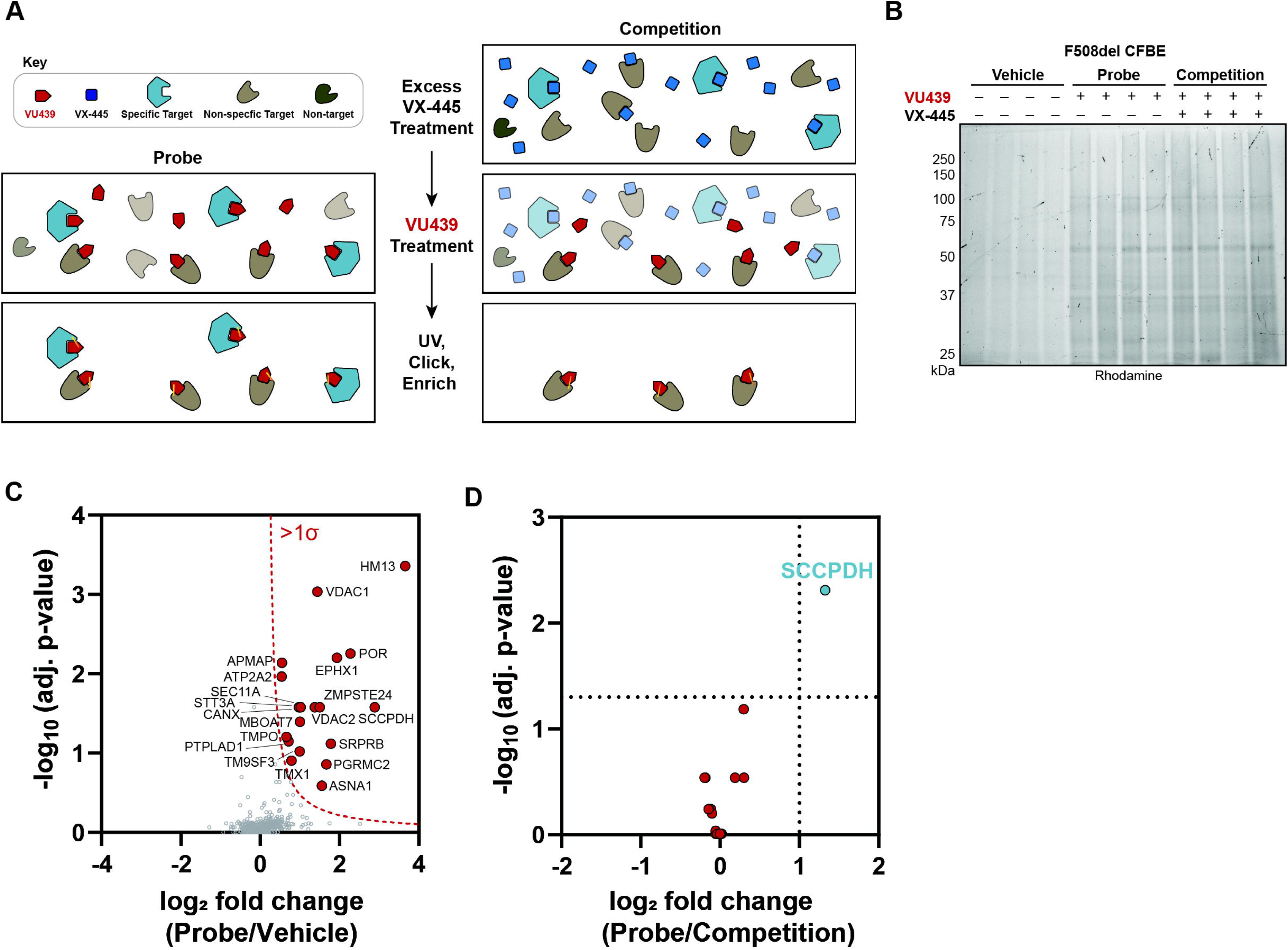
SCCPDH was identified as an off-target of VX-445 via affinity purification mass spectrometry. **A.** Off-target identification workflow. Cells are treated with VU439, UV-crosslinked, and click-modified with TAMRA desthiobiotin azide (TMR). Streptavidin-enriched proteins are digested and analyzed by LC-MS/MS. Competition with parent compound results in loss of specific targets. **B.** Representative SDS-PAGE gel image showing streptavidin-enriched proteins in doxycycline inducible CFBE F508del cells treated with vehicle (DMSO) or PAL probe (VU439 at 1 µM) or competition (VU439 at 1 µM, VX-445 at 10 µM). Addition of probe allows enrichment of crosslinked proteins modified by TMR. **C.** Workflow A was followed and analyzed by data independent acquisition (DIA) mass spectrometry (n = 8). Volcano plot shows log_2_ fold change of enriched protein abundance of probe condition compared to vehicle. Enriched proteins with a standard deviation of at least one from the normal distribution of all identified proteins were selected (red dots) for comparison against the competition condition. **D.** Log_2_ fold changes of filtered proteins were compared to competition conditions to identify probe-specific targets (red). SCCPDH showed clear enrichment effectively competed by parent compound as a specific off-target of VX-445 (blue).

We next performed data independent acquisition-mass spectrometry (DIA-MS) to identify enriched proteins. Protein abundances were normalized to the global median (**Figure S2A, Table S1**). We filtered identified proteins from both probe (VU439: 1 µM) and competition conditions (VU439: 1 µM, VX-445: 10 µM) against vehicle control (DMSO) to gain a list of PAL-dependently enriched targets (**Figure 2C**). We then compared the probe against the competition conditions to remove non-specific targets (**Figure 2D**). Among the twenty proteins enriched compared to background, saccharopine dehydrogenase-like oxidoreductase (SCCPDH) enrichment was lost upon competition with excess VX-445, indicating its specificity as a binding partner. SCCPDH is a 429 amino-acid-long protein with uncharacterized function. Protein and peptide level abundances showed clear enrichment of SCCPDH upon probe addition and loss of enrichment upon competition (**Figure S2B and S2C**).

### II. Validation of SCCPDH as an off-target of VX-445

To further validate SCCPDH as an off-target of VX-445, we transiently overexpressed Myc-tagged SCCPDH in HEK293T cells. Cells were treated with VU439 and irradiated with UV light to capture binding partners. Lysates were modified with the trifunctional probe and crosslinked proteins were enriched with desthiobiotin-streptavidin pulldown. We then performed immunoblotting to probe for SCCPDH-Myc. We observed a dose-dependent increase in enrichment of VU439-TMR labeled proteins including SCCPDH as indicated by both rhodamine and Myc bands (**Figure 3A and 3B**). Addition of VX-445 in five or ten-fold excess decreased enrichment of SCCPDH by 50% or 70% respectively, indicating that excess VX-445 outcompetes VU439 binding to SCCPDH. TMR and Myc bands co-localized at ∼45 kDa confirming direct VU439 labeling of SCCPDH.

**Figure 3.**
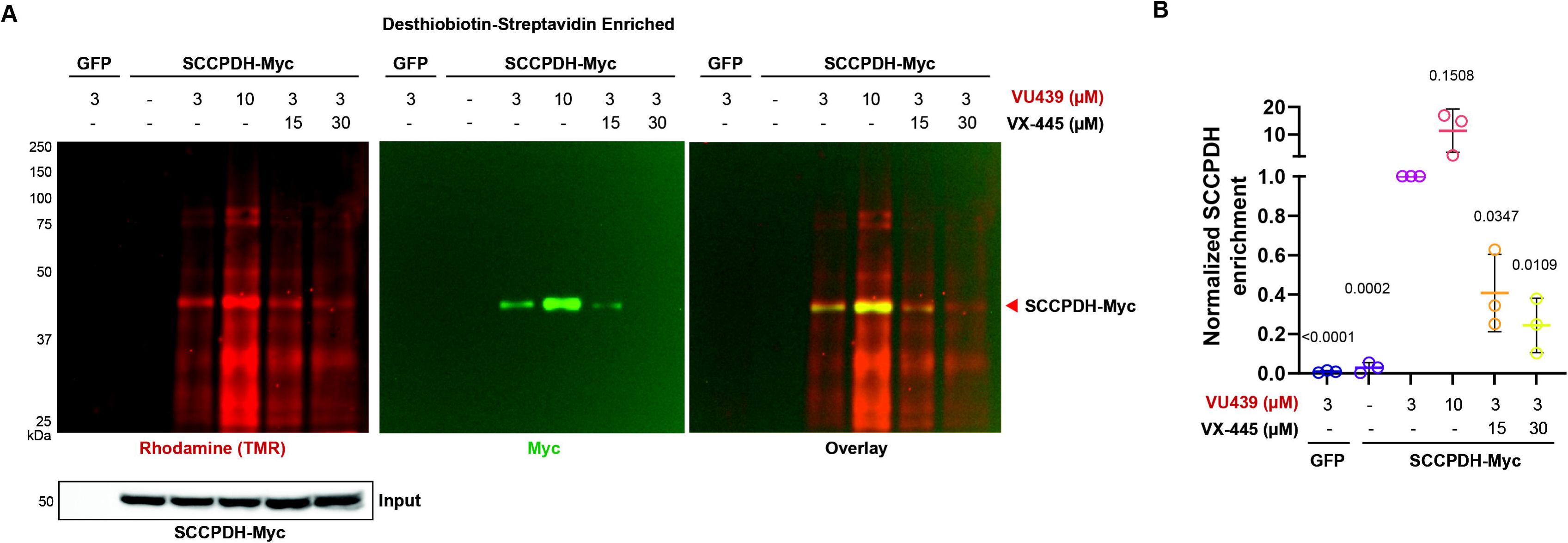
Validation of SCCPDH as an off-target of VX-445. **A.** Representative immunoblot showing enrichment of PAL-modified SCCPDH by desthiobiotin-streptavidin pulldown. HEK293T cells transiently expressing SCCPDH-Myc construct were treated with probe, UV crosslinked and clicked with TMR. SCCPDH-Myc was enriched dose-dependently. Competition with VX-445 at five or ten-fold excess led to dose-dependent loss of enrichment. **B.** Quantification of labeled SCCPDH normalized to 3 µM VU439 treatment reiterates specificity of VU439 binding (n = 3). Statistical differences were computed via one sample t test comparing against 3 µM VU439 treatment. P-values as shown. VX-445 outcompeted VU439 binding to SCCPDH.

### III. Global untargeted metabolomics reveal a role of SCCPDH in amino acid metabolism

We next sought to characterize SCCPDH because its function has yet to be well defined. Sequence homology evidence suggests SCCPDH may be an enzyme involving oxidoreductase activity. To gain insights into the metabolic perturbations elicited by SCCPDH, we performed global untargeted metabolomics. HEK293T cells were transfected with mock (GFP) and treated with DMSO or VX-445, or cells were transfected with SCCPDH to overexpress the putative enzyme (**Figure 4A**). Metabolites were extracted from cellular lysates and analyzed via LC-MS/MS.

**Figure 4.**
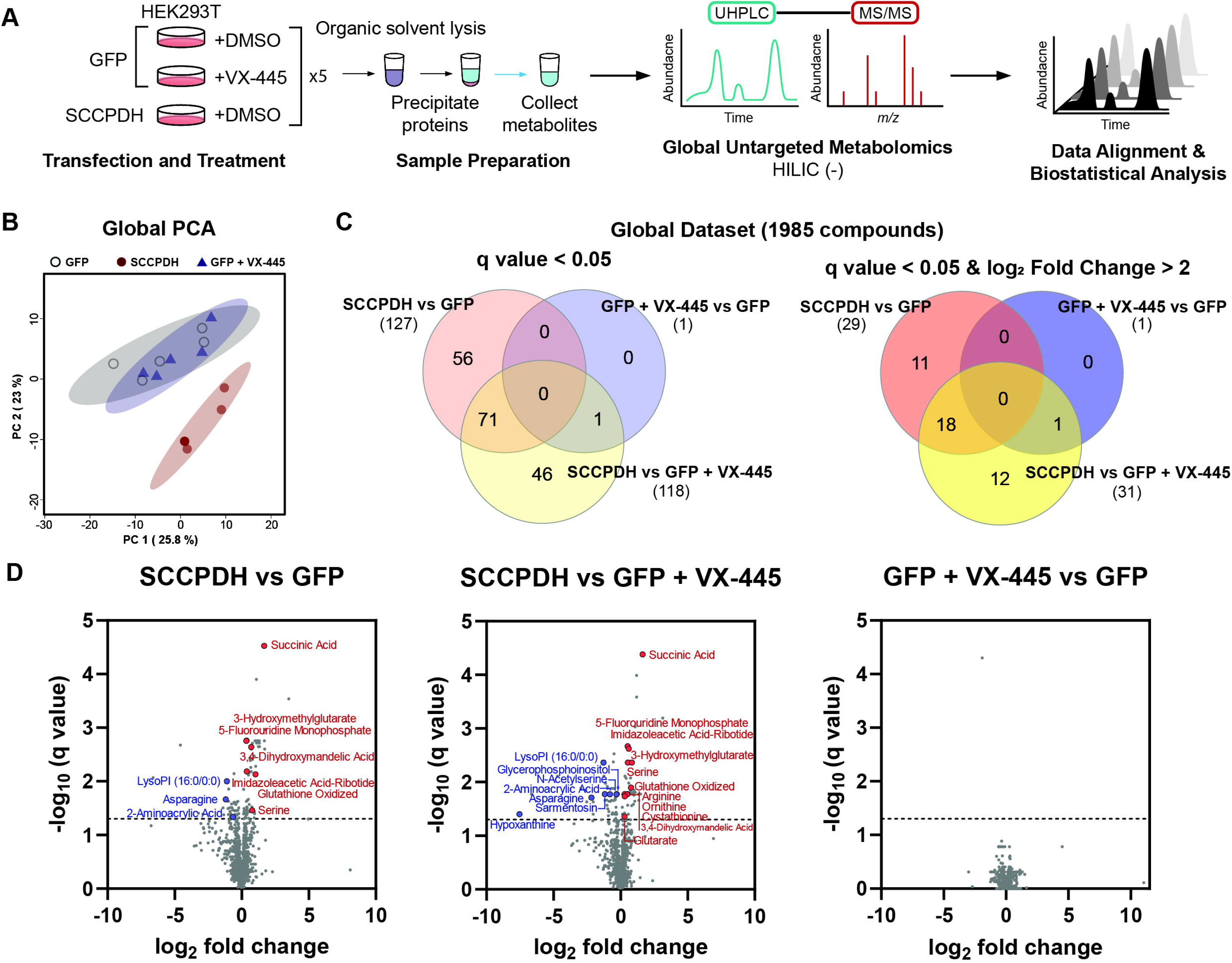
Global untargeted metabolomics reveal a role of SCCPDH in amino acid metabolism. **A.** Schematic of global untargeted metabolomics study. HEK293T cells transfected with GFP or SCCPDH were treated with vehicle (DMSO) or VX-445 (3 µM) (n = 5). Metabolites were extracted from cell pellets for global untargeted metabolomics analysis. UHPLC-MS/MS data was processed and analyzed to identify key metabolites related to SCCPDH. **B.** Global principal components analysis (PCA) plot showing distribution of sample groups as labeled. Samples overexpressing SCCPDH form a cluster separate from GFP +/-VX-445. **C.** Globally, 1985 compounds were detected and have a CV < 25%. Venn diagrams show moderate overlap of compounds identified with FDR < 5% between the comparisons in SCCPDH vs GFP and SCCPDH vs GFP + VX-445. **D.** Volcano plots showing each comparison. Compounds with high confidence (L1, L2) of ID are labeled. See also Table S3.

A clear distinction in metabolome compositions between GFP +/-VX-445 and SCCPDH conditions was observed as shown by the PCA plot (**Figure 4B**). Globally, 1985 compounds were detected in our samples. Comparisons between conditions SCCPDH vs GFP and SCCPDH vs GFP + VX-445 showed 127 and 118 compounds respectively to be significantly altered (FDR q value < 0.05) with 71 compounds overlapping (**Figure 4C, Table S2**). Upon further filtering for log_2_ fold change > 2, we found 29 and 31 compounds for the two comparisons respectively with an overlap of 18 compounds. Comparison between GFP + VX-445 vs GFP showed one compound significantly altered in both filtering methods that intersected with comparison between SCCPDH vs GFP + VX-445.

Compounds or metabolites were annotated using previously established confidence levels (see **Table S3**)^22^. Significantly altered (FDR q value < 0.05) metabolites impacted by SCCPDH overexpression and VX-445 treatment were annotated (19 compounds with L1 and L2 annotations). Overexpression of SCCPDH compared to GFP led to increases in abundance for several metabolites: succinic acid, 3-hydroxymethylglutarate, 5-fluorouridine monophosphate, 3,4-dihydroxymandelic acid, imidazoleacetic acid-ribotide, glutathione oxidized, and serine (**Figure 4D**). These data also show decreased abundance in LysoPI (16:0/0:0), asparagine, and 2-aminoacrylic acid. When SCCPDH overexpression condition was compared to GFP + VX-445, additional metabolites such as arginine, ornithine, cystathionine, and glutarate were increased whereas glycerophosphoinositol, N-acetylserine, sarmentosin, and hypoxanthine were decreased highlighting the impact of VX-445 on endogenously expressed SCCPDH.

### IV. Pathways change with SCCPDH overexpression and VX-445 drives these changes in the opposite direction

To identify metabolic pathways altered by SCCPDH and VX-445, the 19 metabolites (L1 and L2 annotations) with significant changes were analyzed. Heatmap shows hierarchically clustered metabolites of interest (**Figure 5A and S3**). We observed two large clusters in which metabolites were increased or decreased in abundance upon SCCPDH overexpression. Within these clusters, several metabolites (red and blue boxes in **Fig. 5A**) displayed opposing fold changes between the SCCPDH overexpression conditions and the VX-445 treatment. These metabolites were particularly interesting, as VX-445 may act on the endogenous SCCPDH population to drive these metabolic changes.

**Figure 5.**
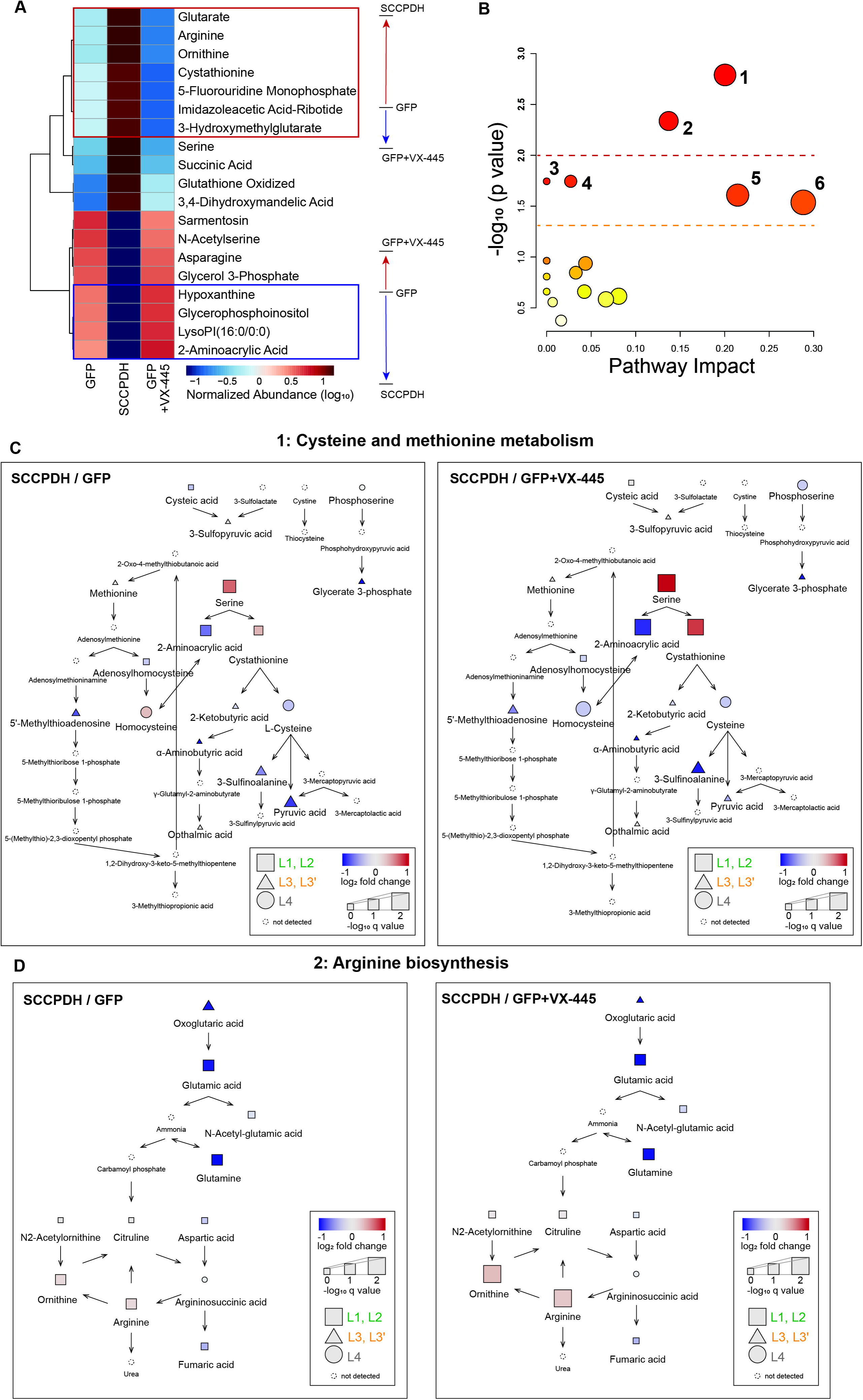
Pathways change with SCCPDH overexpression and VX-445 drives these changes in the opposite direction. **A.** Heatmap shows metabolites significantly altered across all conditions (FDR < 5%, L1 and L2 annotation only). Metabolites were hierarchically clustered using Pearson distance and average clustering. Among the metabolites that drastically change with SCCPDH overexpression, VX-445 treatment further enhanced the observed changes. See also Figure S3. **B.** Pathway analysis performed with the 19 metabolites from A. Cysteine and methionine metabolism (1), and arginine biosynthesis (2) were the most significant at p < 0.01. Alanine, aspartate and glutamate metabolism (3), glutathione metabolism (4), glycine, serine and threonine metabolism (5), and arginine and proline metabolism (6) were observed at p < 0.05. Other pathways at p > 0.05 are listed in Table S4 in order of decreasing significance. Node color and size represent p value and pathway impact score, respectively. **C.** Cysteine and methionine metabolic pathway (1) from B. Overexpression of SCCPDH resulted in serine accumulation and decrease of downstream metabolite 2-Aminoacrylic acid. Addition of VX-445 in GFP control further highlighted this finding. Metabolites detected in our dataset were filled into the pathway as annotated in the key. See also Figure S4. **D.** Arginine biosynthesis pathway (2) from B. Overexpression of SCCPDH resulted in loss of glutamic acid and glutamine with a small increase in ornithine and arginine. Addition of VX-445 in GFP control further highlighted this finding. Metabolites detected in our dataset were filled into the pathway as annotated in the key. See also Figure S5.

Canonical metabolomic pathway analysis of the 19 metabolites (L1 and L2 metabolites with significant changes in abundance) highlight two metabolic pathways significantly altered at p < 0.01: 1) cysteine and methionine metabolism, and 2) arginine biosynthesis (**Figure 5B**). Four other pathways at p < 0.05 were 3) alanine, aspartate and glutamate metabolism, 4) glutathione metabolism, 5) glycine, serine and threonine metabolism, and 6) arginine and proline metabolism. A full list of metabolic pathways altered, statistical metrics, and components is reported in **Table S4**.

We populated the two most significant pathways with metabolites detected in our dataset to understand metabolic dysregulations caused by SCCPDH overexpression and VX-445 treatment. First, in the cysteine and methionine metabolism pathway, overexpression of SCCPDH resulted in accumulation of serine and depletion of its downstream metabolite 2-aminoacrylic acid compared to GFP (**Figure 5C and S4**). Cystathionine, another downstream metabolite of serine, was instead increased, indicating preferential breakdown of serine towards cystathionine in the presence of excess SCCPDH. Moreover, metabolites downstream of cystathionine showed decreasing trends, indicating accumulation of cystathionine. When comparing SCCPDH versus GFP treated with VX-445, the fold changes of these metabolites were further increased, supporting evidence that VX-445 may inhibit endogenous SCCPDH, leading to the opposite effect to that caused by SCCPDH overexpression.

We next investigated the arginine biosynthesis pathway where overexpression of SCCPDH caused increased abundance of arginine and ornithine but decreased abundance of oxoglutaric acid, glutamic acid and glutamine compared to GFP (**Figure 5D and S5**). These data suggest that the overexpression of SCCPDH induces depletion of upstream metabolites such as glutamic acid in favor of the production of downstream metabolites such as arginine. These trends were intensified when comparing SCCPDH versus GFP treated with VX-445, similar to the effect observed in the cysteine and methionine metabolism pathway. These data together support the role of SCCPDH in amino acid metabolism and the potential inhibitory effect of VX-445 on SCCPDH.

We next examined the saccharopine pathway due to the homology profile of SCCPDH to saccharopine dehydrogenase (SDH) which metabolizes saccharopine into α-aminoadipate-δ-semialdehyde and glutamate. In our dataset, among the metabolites involved in the saccharopine pathway, we detected lysine, oxoglutaric acid, saccharopine, glutamic acid, and α-aminoadipate. Interestingly, saccharopine was significantly increased in abundance with SCCPDH overexpression while upstream and downstream metabolites oxoglutaric acid, glutamic acid, and α-aminoadipate were significantly decreased (**Figure S6A and S6B**).

To make sure the most significantly different metabolite based on p value and fold change, succinic acid, was not overlooked, we examined the TCA cycle. Several TCA cycle components such as citrate, fumarate, succinic acid and oxoglutaric acid were detected and identified with high confidence in the dataset. Among these metabolites, succinic acid was significantly increased in abundance with SCCPDH overexpression whereas oxoglutaric acid and fumarate were significantly decreased in abundance with SCCPDH overexpression (**Figure S6C and S6D**). Interestingly, oxoglutaric acid is involved in both the saccharopine pathway and the TCA cycle as precursors to both saccharopine and succinate. Taken together, SCCPDH overexpression results in the conversion of oxoglutaric acid into a buildup of succinate and saccharopine, leading to decreased abundance of the downstream metabolites including glutamate which may be implicated in psychological symptoms observed in CF patients.

Lastly, we examined global proteomes in HEK293T cells to identify global proteomic changes upon SCCPDH overexpression. HEK293T cells expressed ∼32 fold more SCCPDH upon transfection compared to mock expressing endogenous SCCPDH (**Figure S7A, Table S5**). CFBE TetON P67L cells expressed a slightly lower amount of endogenous SCCPDH compared to HEK293T cells. We observed no statistically significant global changes in protein expression caused by SCCPDH overexpression indicating that it is unlikely that the metabolome perturbations are due to wider proteome changes (**Figure S7B**).

## Discussion

Here we report an off-target identification study of the CF corrector VX-445. The synthesized PAL probe retained mild correction of F508del CFTR trafficking and function. Through chemoproteomics, SCCPDH was found to be a specific off-target of VX-445. In these data, we also observed several other targets such as VDAC1, which has been reported previously^23^. These were not outcompeted by a ten-fold excess treatment of VX-445, indicating their lack of specificity to VX-445. Interestingly, SCCPDH was reported by two other groups as specific off-targets of two different small molecules WOBE437^20^ and AX-1^21^. These groups both utilized alkyl diazirine and terminal alkyne modified probes identical to our study. WOBE437 imitates anandamide, a signaling lipid, whereas AX-1 is an AXL kinase inhibitor highlighting the possibility of highly promiscuous binding of SCCPDH to these small molecules.

SCCPDH is not fully annotated and gene ontology search suggests its involvement in lipid metabolism inferred from an ancestral gene. Interestingly, in proximity labeling proteomics studies, SCCPDH was found enriched in lipid droplets, suggestive of its role in redox metabolism in lipid droplets^26,27^. We note that SCCPDH contains a PDZ motif at the C-terminus which is recognized by membrane scaffolding proteins such as GRIP to facilitate its localization which may explain its localization to lipid droplets or potentially transport vesicles at the synapse^28^. CFTR also contains a PDZ motif^29^, and the question remains whether CFTR can interact with SCCPDH via PDZ proteins. Yet, gene knockdown of SCCPDH by RNAi in macrophages was reported to show no significant perturbations in lipid metabolism^30^. Alternatively, in a comparative proteomics study, SCCPDH, along with six other proteins has been shown to be gradually down-regulated after intranigral (within the substantia nigra) grafting in a Parkinson’s disease mice model^31^. Authors suggest that these proteins are important for lipid formation and dopamine vesicle recycling at the synapse. Furthermore, in another proteomics study, SCCPDH was found upregulated in anxiety-susceptible rat under chronic stress^32^. These findings suggest a potential role of SCCPDH in dopamine regulation that may act as an underlying cause of psychological symptoms such as depression and anxiety. These symptoms are commonly observed in CF patients at baseline and approximately 10% of patients reported worsening psychological symptoms and sleep disorders after starting ETI^33–37^. Disruption of SCCPDH function by VX-445, a component of ETI, could possibly be associated with this outcome (**Figure 6**). Other disease contexts related to SCCPDH include transcriptomic dysregulation in ischemia, and as a diagnostic marker for preeclampsia^38,39^.

**Figure 6.**
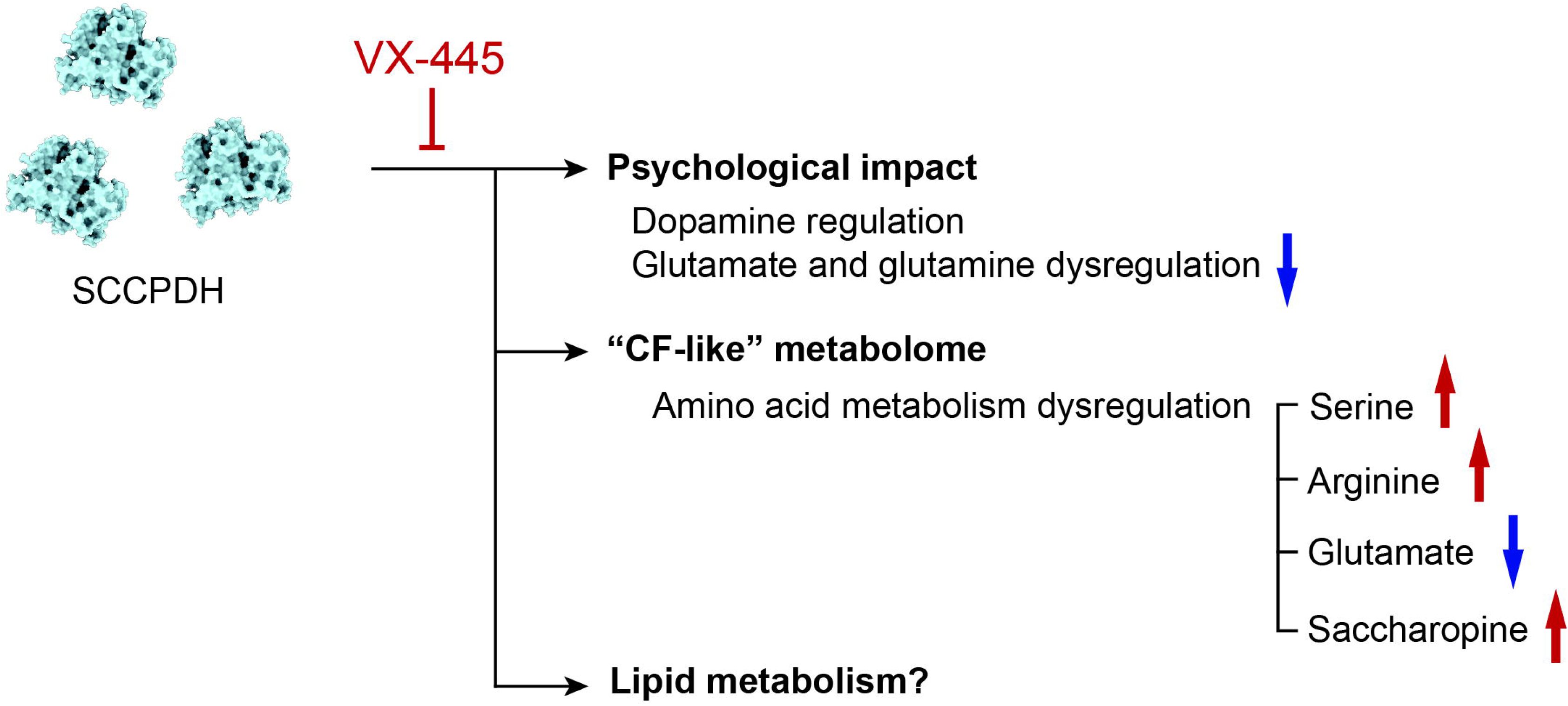
Inhibition of SCCPDH by VX-445 may result in amino acid metabolism dysregulation resulting in side effects. Amino acids such as glutamate and glutamine were decreased upon overexpression of SCCPDH and VX-445 mitigated this decrease which may be the underlying cause of psychological symptoms experienced by patients on VX-445. Additionally, metabolites reported to be dysregulated in CF were observed to change in identical directions upon SCCPDH overexpression. VX-445 may inhibit SCCPDH to counteract these dysregulations resulting in unforeseen outcomes such as side effects.

In our study, SCCPDH overexpression led to increased amounts of saccharopine, which was consistent with a decrease in its precursor oxoglutaric acid (or α-ketoglutarate) but its other precursor lysine remained unchanged (**Figure S6A and S6B**). We next noted that succinic acid (or succinate) another downstream metabolite of oxoglutaric acid increased significantly with SCCPDH overexpression (**Figure S6C and S6D**). Thus, we attribute the decrease in oxoglutaric acid to the conversion into succinate rather than into saccharopine. Consequently, the accumulation of saccharopine points to decreased breakdown rather than increased production, resulting in decreased glutamate production. We speculate a possible binding and sequestration of saccharopine by SCCPDH rendering it inaccessible for metabolism by SDH.

Metabolic pathway analysis showed decreased glutamic acid, consistent with the potential outcomes of dysregulation in both the saccharopine pathway and the TCA cycle. Glutamate has many important functions as a neurotransmitter and a precursor to other amino acids which may drive psychological conditions in its absence. Moreover, several amino acids changed with SCCPDH overexpression. It is unclear whether SCCPDH directly regulates these amino acids, or it indirectly causes a shift in amino acid metabolism as a result of glutamate dysregulation. Similarly, there may be connections to phenylalanine and tyrosine metabolism, which cells undergo to generate dopamine.

Metabolomics studies involving CF have been largely focused on biomarker discovery in various patient samples^40–44^. In a first-in-line metabolomics study, primary human airway epithelial cell cultures from CF patients exhibited differences in nucleotide metabolism, amino acid metabolism, glutathione, osmolytes and glucose metabolism^45^. In these pathways, CF patients showed decreases in metabolites such as hypoxanthine, nicotinamide, glucose, and fructose, indicating suppression of these pathways. Similarly, glucose and amino acid metabolism was shown to be dysregulated in CF patient blood samples^46^. Interestingly, we found differentially expressed metabolites upon SCCPDH overexpression to overlap and trend similarly with those found to be dysregulated in CF patients such as glutamic acid, arginine, hypoxanthine, saccharopine, oxoglutaric acid, and ornithine. The presence of excess SCCPDH therefore appears to drive the cellular metabolic state towards a “CF-like” state. We observed treatment of VX-445 neutralizes this “CF-like” state in the absence of CFTR in HEK293T cells. This suggests an alternative mechanism of action by VX-445 to normalize cellular metabolic state by acting on SCCPDH.

This study is limited to surveying the CFBE cell model for off-targets. Surveying other CF-relevant cell lines such as neuronal, lung, and intestinal may provide a better scope of binding partners. Systemic side-effects stemming from VX-445 binding to SCCPDH are difficult to predict. The synthesized probe was limited to copper-mediated click chemistry, which leaves membrane proteins prone to aggregation during the precipitation clean-up step. This renders membrane proteins insoluble and subsequently undetectable by MS analysis. In addition, performing lipidomic studies to further characterize SCCPDH may reveal a connection to CFTR, since lipids such as cholesterol are highly implicated in CFTR trafficking and function^44,47,48^.

In summary, we report a functionalized PAL for CF corrector VX-445 and show off-target identification via LC-MS/MS that can be applied to other pharmaceuticals to identify their target spectrum. We identified SCCPDH as an off-target for VX-445. Metabolomics investigation of overexpressed SCCPDH in cells showed changes in amino acid metabolic pathways consistent with CF-induced changes. VX-445 drove these changes in reverse even in the absence of CFTR, suggesting its inhibitory function on SCCPDH through direct binding rather than through a secondary effect from correction of CFTR. These findings may provide insights into the indirect modes of clinical efficacy or explain adverse effects observed in patients. Further optimization of correctors in consideration of their target spectrum will therefore benefit the subpopulation of patients who are intolerant to existing therapies. VX-121 (Vanzacaftor), a cyclized structural analog of VX-445 was approved by the FDA in December 2024, replacing VX-445 in ETI^49^. It remains to be observed whether side effects are diminished compared to VX-445.

## Resource Availability

The mass spectrometry proteomics data have been deposited to the ProteomeXchange Consortium via the PRIDE^50^ partner repository with the dataset identifier PXD060898. The untargeted metabolomics data is available at the NIH Common Fund’s National Metabolomics Data Repository (NMDR) website, the Metabolomics Workbench, https://www.metabolomicsworkbench.org, under the assigned Study ID ST003713. The data can be accessed directly via its Project DOI: http://dx.doi.org/10.21228/M8JR73. This resource was supported by NIH grants U2C-DK119886 and OT2-OD030544.

## Acknowledgments

This work was supported by the National Institute of General Medicine Sciences (R35 GM133552); National Heart, Lung, and Blood Institute (R01 HL147351, R01 HL1670146), National Institute of Diabetes and Digestive and Kidney Diseases (P30-DK117467) and Vanderbilt University funds. We thank Dr. Guido Veit and Dr. Gergely Lukacs (McGill University) for sharing CFBE41o-TetON CFTR expressing cells. We thank members of the Plate lab for their critical reading and feedback on this manuscript. VU439 was synthesized by the Vanderbilt Institute of Chemical Biology, Molecular Design and Synthesis Center, Vanderbilt University, Nashville, TN 37232-0412. This work was supported in part using the resources of the Center for Innovative Technology (CIT) at Vanderbilt University. Figure 1A was generated using Biorender.com.

## Author Contributions

Conceptualization (L.P., G.A.S., M.K.); Investigation (M.K.); Data curation & Formal Analysis (M.K., K.K., L.A.B., J.L., S.G.C., S.D.S., I.R., K.J.); Funding acquisition (L.P., A.P.N.); Methodology (M.K., K.K., J.L., L.A.B.), Project administration (L.P., G.A.S., J.A.M., A.P.N.); Writing – original draft (M.K, L.P.); Writing – review & editing (all authors)

## Supplemental Information

Document S1. Figures S1-S7, Tables S3-S4, and Synthesis of Photoaffinity Ligand VU439

Table S1 Excel file containing AP-MS processed data

Table S2 Excel file containing normalized metabolomics data

Table S5 Excel file containing whole cell proteomics processed data

## Experimental Procedures

### Plasmids and Antibodies

Plasmids used for transient transfection expressed WT or F508del CFTR in the pcDNA5/FRT vector, or SCCPDH-Myc (Sino Biological, HG17073-CM) in the pCMV3 vector. Anti-CFTR antibodies used for detection were 217 and 596 (provided by J. Riordan, University of North Carolina, Chapel Hill, North Carolina; http://cftrantibodies.web.unc.edu/) each at 1:1000 and 1:500 working dilutions in immunoblotting buffer (5% bovine serum albumin [BSA] in Tris-buffered saline, pH 7.5, 0.1% Tween-20, and 0.1% NaN_3_), respectively. Primary antibodies used were anti-myc 9B11 (1:1000, mouse monoclonal, Cell Signaling Tech, 2276S), rhodamine–conjugated tubulin (1:10000, Bio-Rad, 12004165). Streptavidin-IRDye 680RD (1:5000, LI COR, NC0883597) was used to probe for biotin. Secondary antibodies used were goat anti-mouse StarbrightB700 (Bio-Rad, 12004158), anti-mouse IgG HRP conjugate (Promega, W4021).

### Cell Culture

Human embryonic kidney 293T (HEK293T) cells were cultured in Dulbecco’s modification of Eagle’s medium (DMEM, Corning) supplemented with 10% fetal bovine serum (FBS, Gibco), 1% L-glutamine (200 mM, Gibco), and 1% penicillin/streptomycin (10,000 U; 10,000 μg/mL, Gibco).

TetON inducible Cystic Fibrosis bronchial epithelial cells (CFBE) were generously gifted by Dr. Guido Veit and Dr. Gergely Lukacs, McGill University^51^. CFBE cells expressing F508del CFTR were cultured in minimum essential medium (MEM, Gibco) supplemented with 10% FBS (Peak), 1% HEPES (1 M, Gibco), and 1% L-glutamine (200 mM, Gibco). For experiments, CFBE cells were allowed to differentiate for at least 72 h after full confluency. Induction of CFTR expression was performed with 500 ng/L doxycycline (Fisher Bioreagents) for at least 72 h before collection. Media was replenished every 48 h.

Cells were maintained in a 37 °C humidified incubator at 5% CO_2_ – 95% air.

### Immunoblotting

HEK293T cells were transiently transfected with CFTR plasmids via calcium-phosphate transfection^52^. Cells were exposed to respective drug treatments at 3 μM for 24 h before collection. Cells were rinsed with ice cold phosphate-buffered saline (PBS) and lysed on plate with 600 μL of TNI buffer (50 mM Tris-base, 150 mM NaCl, pH 7.5, and 0.5% IGEPAL CA-630, EDTA-free protease inhibitor cocktail [Roche]) rocking at 4 °C for 20 min. Cells were harvested by scraping and sonicated for 3 min and centrifuged at 18,000×g for 30 min. Resulting lysate was normalized with BCA protein assay kit (Pierce, 23225) to contain equal amounts of total protein. Samples were denatured in 2.4× Laemmli buffer with 10 mM dithiothreitol, resolved by 8% SDS– PAGE, and transferred to PVDF membranes (Millipore). Membranes were blocked in 5% milk in Tris-buffered saline, pH 7.5, 0.1% Tween-20 (TBS-T) at RT rocking for 30 min. Membranes were washed three times with TBS-T and probed in primary antibodies overnight at 4 °C. After three washes with TBS-T, membranes were probed with secondary antibody in 5% milk TBS-T at RT for 30 min. Membranes were washed three times with TBS-T and imaged using a ChemiDoc MP Imaging System (Bio-Rad). Quantification was performed using ImageLab (Bio-Rad).

### Whole Cell Patch Clamp

HEK293 cells were transfected with WT or F508del CFTR for 24 h using Lipofectamine 3000 (ThermoFisher, Cat#L3000015). F508del CFTR was treated with VX-445 (MedChemExpress, Cat#HY-111772) or VU439 in culture medium for 24 h prior to experiment. Stimulation was performed with forskolin (20 μM) and inhibition by CFTR inh-172 (20 μM). Cells were positively selected for co-transfected GFP (Addgene, Cat#74165). Whole cell patch clamp recordings were acquired using an Axopatch-200B amplifier connected to Axon DigiData 1550B (Molecular Devices, CA, USA). Patch pipettes with resistances of 3 - 6 MΩ were filled with pipette solution and prepared using a micropipette puller (Sutter Instrument, CA, USA, Cat# P-1000). To simultaneously obtain current traces at -60 mV and I/V curves of CFTR, whole-cell currents were consecutively recorded with a 1 s voltage ramp of ± 100 mV applied every 10 s: hold at Vm = -60 mV and filtered at 1 kHz and sampled at 50 Hz. The pipette solution was composed of (in mM): 116 NMDG-Cl^-^, 30 aspartic acid, 1 MgCl_2_, 5 ethylene glycol tetraacetic acid (EGTA), 2.9 CaCl_2_, 10 HEPES, and 3 Mg-ATP, titrated to pH 7.4 with HCl. The bath solution was composed of (in mM): 146 NMDG-Cl^-^, 1 CaCl_2_, 1 MgCl_2_, 10 Glucose, 10 HEPES, titrated to pH 7.4 with HCl^53^.

### Photoaffinity Labeling and Click Chemistry

TetON inducible CFBE cells expressing F508del were seeded at 5×10^5^ cells/mL and induced with 500 ng/L doxycycline upon reaching confluency. Cells were treated at 48 h after confluency with vehicle (DMSO) or probe (VU439) or competition (incubated with VX-445 for 1 h, followed by VU439 addition). After incubating cells for 24 h, cell plates with lids removed were exposed to 365 nm for 2 min for a total of ∼600 mJ. Cells were then rinsed three times with ice cold PBS and lysed on plate with 200 µL of radioimmunoprecipitation assay (RIPA) buffer (50 mM Tris-base, 150 mM NaCl, 0.1% SDS, 1% Triton X-100, 0.5% deoxycholate, EDTA-free protease inhibitor cocktail) rocking at 4 °C for 30 min. Cells were harvested by scraping and centrifuged at 16,000×g for 15 min at 4 °C. Total protein concentration of resulting lysate was measured with a BCA protein assay kit (Pierce, 23225) following manufacturer’s protocol. Samples were diluted to 1 mg/mL for click reaction at a volume of 100 µL with the addition of a master mix (final concentrations: 0.8 mM CuSO_4_, 1.6 mM BTTAA, 5 mM sodium ascorbate, and 100 µM TAMRA Desthiobiotin Azide [Click chemistry tools]) or 200 µM DADPS Biotin Azide [Click chemistry tools] for 1 h at 37 °C shaking at 1000 rpm.

### Desthiobiotin-Streptavidin Enrichment

Pulldown of crosslinked proteins was performed as follows. Click reaction mixture was methanol/chloroform (MeOH/CHCl_3_) precipitated using mass spectrometry grade MeOH, CHCl_3_, and water (3:1:3 ratio) and washed three times with MeOH (500 µL) with 3 min centrifugation (21.1×kg, RT). Protein pellets were then air dried to near dryness and resuspended in 100 μL 6 M Urea 1% SDS in PBS by vortex and sonication in a water bath sonicator until no pellets were visible. Resuspension was diluted with the addition of 1 mL PBS. Streptavidin agarose beads were washed three times with PBS before addition of 60 μL of 1:1 slurry to each sample. Mixture was incubated at RT for 24 h on a head-over-head rotator. Beads were rinsed with 1 mL of 1% SDS in PBS, 4 M Urea, 1 M NaCl, and 1% SDS in PBS, then a 30G×1/2 needle and a 1 mL Henke-Ject Luer syringe was used to completely remove liquid. Beads were then frozen at –80 °C for at least 1 h. Proteins were then eluted at 95 °C for 5 min with 60 μL of elution buffer (50 mM biotin, 1% SDS in PBS, pH 7.2). Elution steps were repeated once and combined. A fraction of the elution was denatured with 1× Laemmli buffer with 10 mM dithiothreitol, heated at 95 °C for 5 min to resolve by SDS-PAGE. Gel was imaged using rhodamine channel on a ChemiDoc MP Imaging System (Bio-Rad) to confirm enrichment of crosslinked proteins.

### Label Free DIA Sample Preparation

MS sample preparation of enriched samples was performed as follows. Briefly, samples were precipitated in MeOH/CHCl_3_. The precipitated pellet was rinsed, air-dried, and reconstituted in 3 μL of 1% Rapigest SF in water (Waters, #186002122) via vortex and sonication. Resuspended proteins were subsequently diluted with 34.5 μL of water, 10 μL of 0.5 M HEPES (pH 8.0), 0.5 μL of fresh 0.5 M tris(2-carboxyethyl)phosphine (TCEP, Sigma), 1 μL of fresh 0.5 M iodoacetamide (IAA, Sigma) and digested with 0.5 μg of Trypsin/Lys-C (Thermo Fisher # A40007) for 14 h. After digestion, formic acid (FA; ThermoFisher Scientific, PI28905) was added to each sample to a final concentration of 2% (v/v, pH ∼2) and incubated at 37 °C for 30 min to cleave Rapigest SF. Samples were then centrifuged at 21.1 kg for 15 min RT to transfer supernatant to a fresh low-bind tube (ThermoFisher Scientific, 21-402-902). Samples were subsequently reduced to dryness via SpeedVac and stored at -80 °C until use. Peptides were resuspended in buffer A (4.9% ACN, 95% H_2_0, 0.1% FA (v/v/v) with an additional 1 µL of FA added prior to LC-MS/MS analysis. The sample was then centrifuged at 21.1×kg, RT for 15 min to transfer supernatant into MS autosampler vials.

### DIA LC-MS/MS Analysis

LC-MS/MS analysis was performed with an Exploris480 mass spectrometer (Thermo Fisher) equipped with an Ultimate3000 RSLCnano system (Thermo Fisher). Peptides were separated using a fused silica microcapillary column (ID 100 μm) ending with a laser-pulled tip filled with 21.5 cm of Aqua C18, 3 μm, 100 Å resin (Phenomenex # 04A-4311). Electrospray ionization was performed directly from the analytical column by applying a voltage of 2.2 kV (positive ionization mode) with an MS inlet capillary temperature of 275 °C and a RF Lens of 40%. Sample was loaded onto a commercial trap column (C18, 5 μm, 0.3 × 5 mm; ThermoFisher Scientific, 160454) using an autosampler. Peptides were then eluted and separated on a 2 h gradient with a constant flow rate of 500 nL/min: 2% B (5 min hold) was ramped to 35% B over 90 min and stepped to 80% in 5 min and held at 80% B for 5 min, followed by a 2 min step down and 13 min hold at 4% B to re-equilibrate the column. Between injections, a 45 min column wash was performed using the following gradient: 2% B (6 min hold) stepped to 5% over 2 min and subsequently ramped to a mobile phase concentration of 35% B over 7 min, ramped to 65% B over 5 min, held at 85% B for 8 min, then returned to 3% B for the remainder of the analysis.

MS/MS scans were acquired in a staggered window pattern. Briefly, 62 × 10 m/z (380-1000 m/z) precursor isolation window DIA spectra (30,000 resolution, automatic gain control target 1e6, maximum ion injection time 55 ms, 27 HCD collision energy) were acquired within a single duty cycle to achieve an effective isolation window size of 10 m/z. A precursor spectrum (380-1000 m/z, 120,000 resolution, automatic gain control target 3E5, maximum injection time 240 ms) was collected prior to the acquisition of MS/MS scans across the mass range.

### Peptide Identification and Quantification

DIA spectra were analyzed using DIA-NN (1.8.1). For precursor ion generation, deep learning-based library free search was utilized. The generated spectral library was built using a Uniprot Mus Musculus database (UP000000589). N-terminal M excision and C carbamidomethylation were included as fixed modifications. Peptides 6 to 30 residues in length with one missed trypsin cleavage were included. Precursors between m/z 380-1000 with charge states of 1-4 and fragment ions between m/z 200 and 1800 were included. Smart profiling was used for library generation.

The algorithm was run under the following DIA-NN GUI selections: unrelated runs, match between runs (MBR), heuristic protein inference, and no shared spectra. Protein inference was made on genes, robust LC (high precision) was used as the quantification strategy, and normalization was set to off.

Data was processed with a custom R code (https://github.com/Plate-Lab/main/blob/main/rcode/DIA.R) to generate protein abundances. The proteomics data (protein IDs and quantification) is included in Table S1 and S5.

### Target Identification and Statistical Analysis

Protein abundance values were global median normalized and log_2_ transformed to perform multiple paired t tests with two-stage step-up (Benjamini, Krieger, and Yekutieli) for targets below 5% false discovery rate (FDR). Targets that were enriched over one standard deviation (σ) from the log2 distribution when probe was compared to vehicle were selected for comparison against probe vs competition. Briefly, the histogram of log_2_ fold changes of probe over vehicle were fit to a Gaussian curve using a nonlinear least-square fit to determine σ. Fold change cutoff for interactors was set to 1 σ. A reciprocal curve with the equation y > c/(x − x_0_), where y = p-value, x = log_2_ fold change, x_0_ = fold change cutoff (1 σ), and c = the curvature (c = 0.8) was used to filter for enriched targets. Identical statistical testing was performed on the enriched proteins to yield PAL specific targets with a cutoff of FDR < 5% and log_2_ fold change > 1.

### Global Proteomics

HEK293T cells were transfected with mock or SCCPDH plasmid and seeded in 6-well plates to be harvested at confluency. CFBE cells were seeded in 6-well plates and induced with 500 ng/L doxycycline upon reaching confluency, then harvested after 72 h. Cells were lysed in TNI or RIPA buffer and protein concentration of whole cell lysates was measured using a BCA assay kit (Pierce, 23225) following manufacturer’s protocol. Samples were normalized to 20 µg of total protein. MS samples were prepared following **Label Free DIA Sample Preparation**. Samples were analyzed via **DIA LC-MS/MS Analysis** where 600 ng of protein was injected. Data analysis was done following **Peptide Identification and Quantification**.

### Global Untargeted Metabolomics

HEK293T cells transiently transfected with GFP were either treated with DMSO or VX-445 (3 µM) or cells were transfected with SCCPDH-Myc and treated with DMSO for 24 h (n = 5). Cells were scraped in ice cold ammonium formate (50 mM) and centrifuged at 200×g for 3 min at 4 °C. Cell pellets were frozen at -80 °C until further processing.

Samples were thawed on ice and lysed in 300 µL ice-cold lysis buffer (1:1:2, acetonitrile: methanol: ammonium bicarbonate 0.1M, pH 8.0) followed by probe tip sonication with 10 pulses at 30% power. Protein content was determined using a bicinchoninic acid protein assay (BCA assay, Thermo Fisher Scientific, Waltham, MA) and the appropriate amount of lysate was taken for 200 µg total protein per sample and adjusted to 200 µL total volume with lysate buffer. Isotopically labeled standards (phenylalanine and biotin) were added to each sample to determine sample process variability as previously described^54–57^.

Following volume adjustment to 200 µL, 800 µL of cold MeOH was added to the samples. Individual samples were vortexed for 30 seconds and incubated overnight at -80 °C for protein precipitation. Following incubation, samples were centrifuged for 10 min at 10,000 rpm at 4 °C and the supernatant was transferred to a new labeled tube and dried down using a cold vacuum centrifuge.

Samples were reconstituted in 100 µL H_2_O, 100 µL MeOH, and 10 µL of SPLASH LIPIDOMIX with vortex mixing after each addition. Samples were incubated at room temperature for 10 min followed by liquid-liquid extraction. For liquid-liquid extraction (LLE), 600 µL MTBE was added with vortex mixing for 30 seconds followed by incubation on ice for 10 min and centrifugation at 15,000 rpm for 15 minutes at 4 °C. An upper (hydrophobic) fraction was transferred and dried down using cold vacuum centrifuge and stored at -80 °C for further lipidomic studies. The lower (hydrophilic) fraction was transferred into a new Eppendorf tube, dried *in vacuo*, and stored at -80 °C until further use.

Prior to mass spectrometry analysis, individual hydrophilic extracts were reconstituted in 80 µL acetonitrile/water (80:20, v/v) containing isotopically labeled standards, tryptophan, inosine, valine, and carnitine, and centrifuged for 5 min at 10,000 rpm to remove insoluble material. A pooled quality control (QC) sample was prepared by pooling equal volumes of individual samples following reconstitution. The QC sample allowed for column conditioning (eight injections), retention time alignment, and assessment of mass spectrometry instrument reproducibility throughout the sample set.

LC-MS and LC-MS/MS analyses were performed on a high-resolution Q-Exactive HF hybrid quadrupole-Orbitrap mass spectrometer (Thermo Fisher Scientific, Bremen, Germany) equipped with a Vanquish UHPLC binary system (Thermo Fisher Scientific, Bremen, Germany). Extracts (8 μL injection volume) were separated on an ACQUITY UPLC BEH Amide HILIC 1.7 μm, 2.1 × 100 mm column (Waters Corporation, Milford, MA) held at 30 °C using the LC method as previously described^54,58–61^. Full MS analyses were acquired over the mass-to-charge ratio (*m/z*) range of 70 - 1,050 in negative ion mode. The full mass scan was acquired at 120K resolutions with a scan rate of 3.5 Hz and an automatic gain control (AGC) target of 1×10^6^. Tandem MS spectra were collected at 15 K resolution, AGC target of 2×10^5^ ions, and maximum ion injection time of 100 ms.

Progenesis QI v.3.0 (Non-linear Dynamics, Newcastle, UK) was used to review, process and normalize the mass spectrometry data. The pooled QC sample was used to align all MS and MS/MS sample runs. Unique ions (retention time and m/z pairs) were de-adducted and de-isotoped to generate unique “features” (retention time and m/z pairs). Data were normalized to all features detected and further curated by applying QA practices to the data. Specifically, compounds or metabolites with spectral features > 25% coefficient of variation (CV) in the pooled QC samples were removed. Sample process and instrument variability were also assessed using the normalized measurements of the isotopically labeled standards to determine sample and batch acceptance. QA metrics for sample process variability and instrument variability were ≤ 20% CV and ≤ 10% CV, respectively. Accurate mass measurements (< 5 ppm error), isotope distribution similarity, and fragmentation spectrum matching (when applicable) were used to determine tentative, putative and validated (Level 1-3) annotations^22^. Compounds were searched using Human Metabolome Database (HMDB)^62^ and a highly curated in-house library available in the Center for Innovative Technology at Vanderbilt University. Metaboanalyst 5.0 (www.metaboanalyst.ca/) was used to perform pathway and metabolite enrichment analyses from annotated compounds with statistical significance^63^. The metabolomics dataset (metabolite IDs and quantification) is included in Table S2.

### Synthesis of Photoaffinity Ligand VU439

See supplementary information for full synthetic scheme and characterization.

