## Supplemental Information for "Photoaffinity ligand of Cystic Fibrosis corrector VX-445 identifies SCCPDH as an off-target"

For

| | |
|---|---|
| Figures S1-S7 | p S2-S8 |
| Tables S3-S4 | p S9 |
| Synthesis of Photoaffinity Ligand VU439 | p S10-S16 |
| References | p S17 |

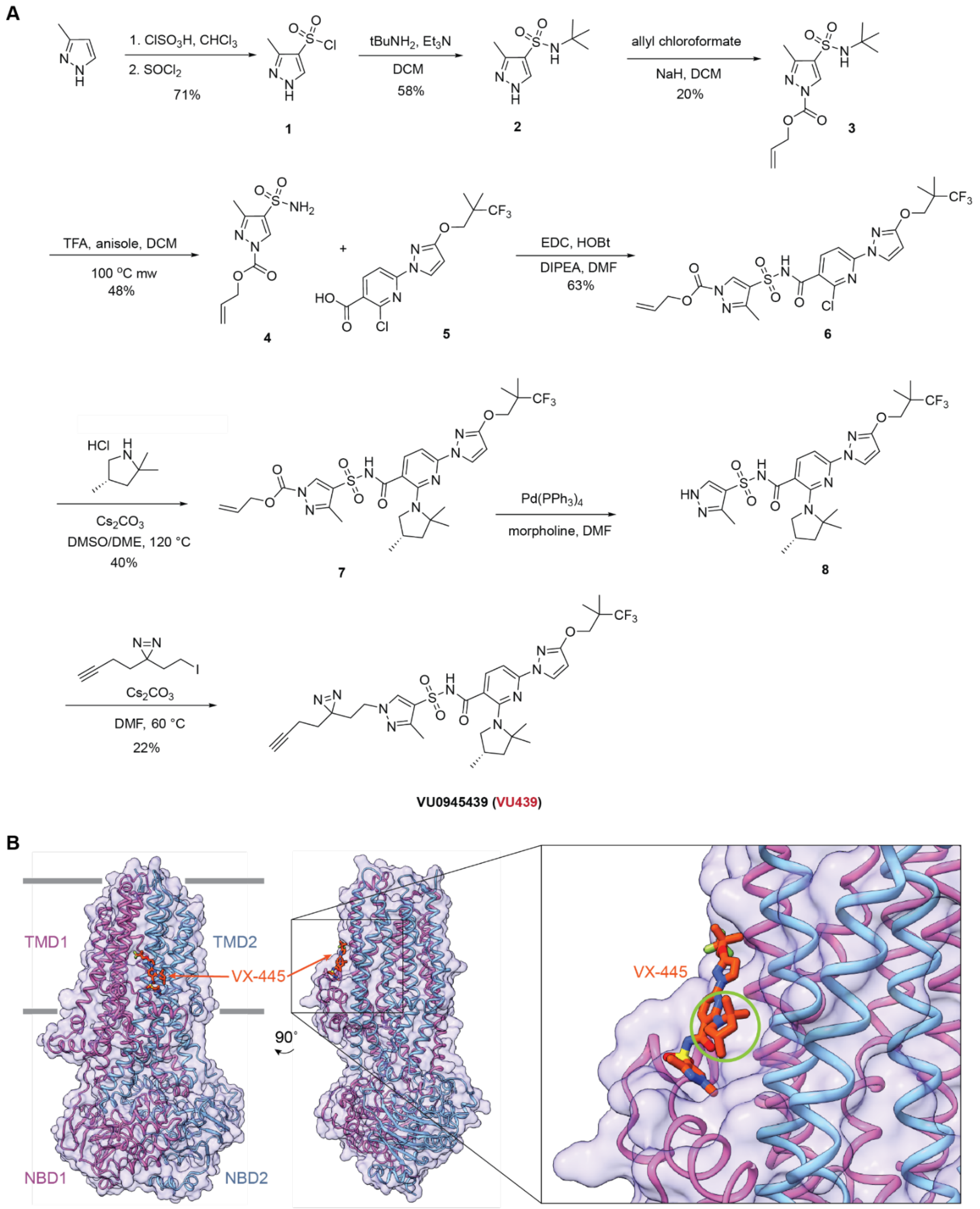


**Figure S1. Synthesis of VU439**

**A.** VU439 was synthesized by the above scheme. Using a newly developed convergent synthesis route, the N-methyl group on the pyrazole was modified to contain a minimalist alkyl diazirine and a terminal alkyne. Detailed conditions and characterization of products are described in the SI.

**B.** VX-445 bound cryo-EM structure (PDB: 8EIQ[1]). Insert shows solvent exposure of pyrazole ring circled in green.

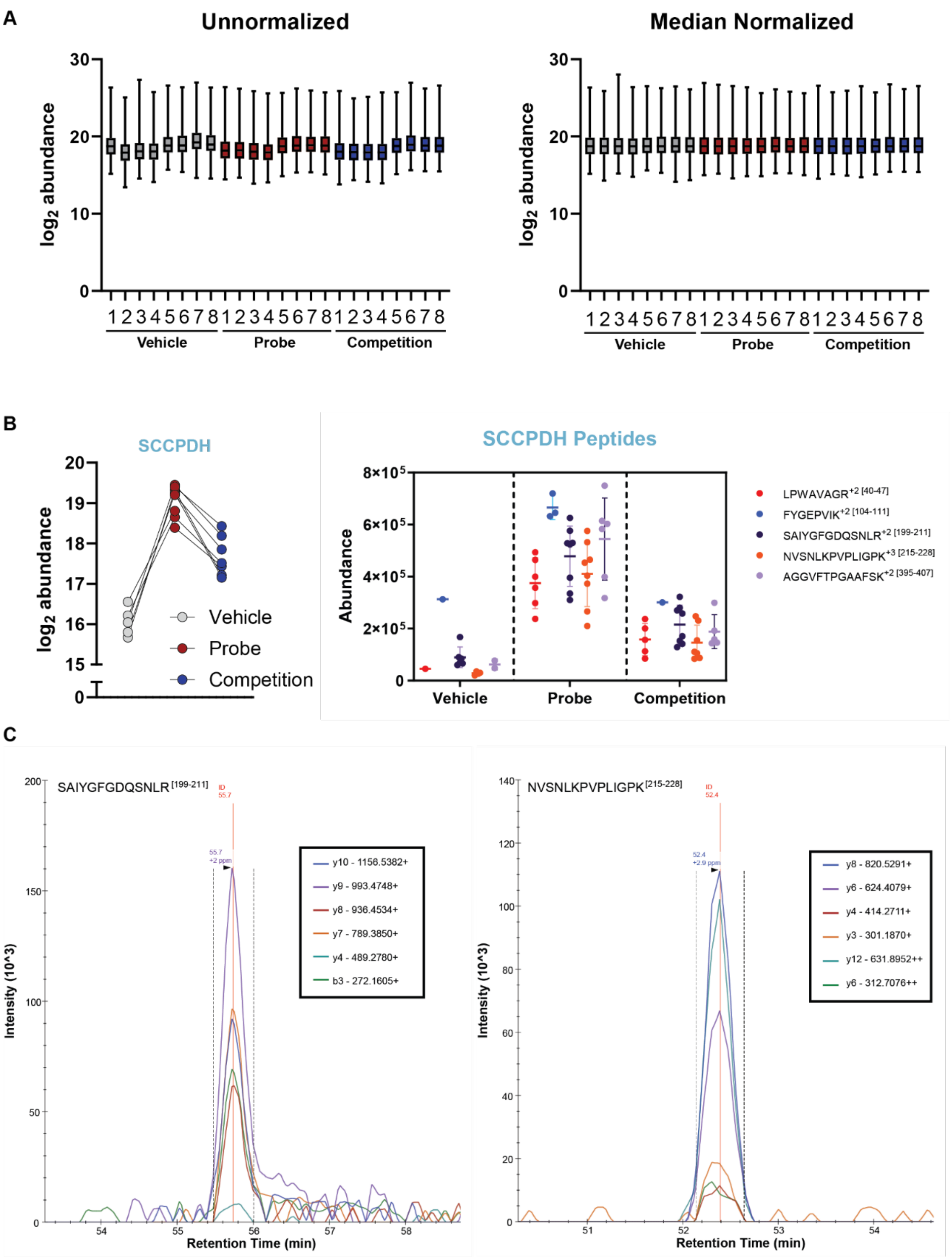


**Figure S2. Validation of VU439 crosslinking and LC-MS/MS analysis.**

**A.** Box plots showing protein abundance distribution for each replicate analyzed via DIA-MS. Abundances were normalized to the global median prior to comparison.

**B.** Protein and peptide level abundances of SCCPDH in each condition shows its enrichment upon probe addition and loss of enrichment upon competition. Each point represents replicates.

**C.** Representative DIA-MS spectra of identified SCCPDH peptides in the probe condition. Figure generated using Skyline (version 23.1.0.268).

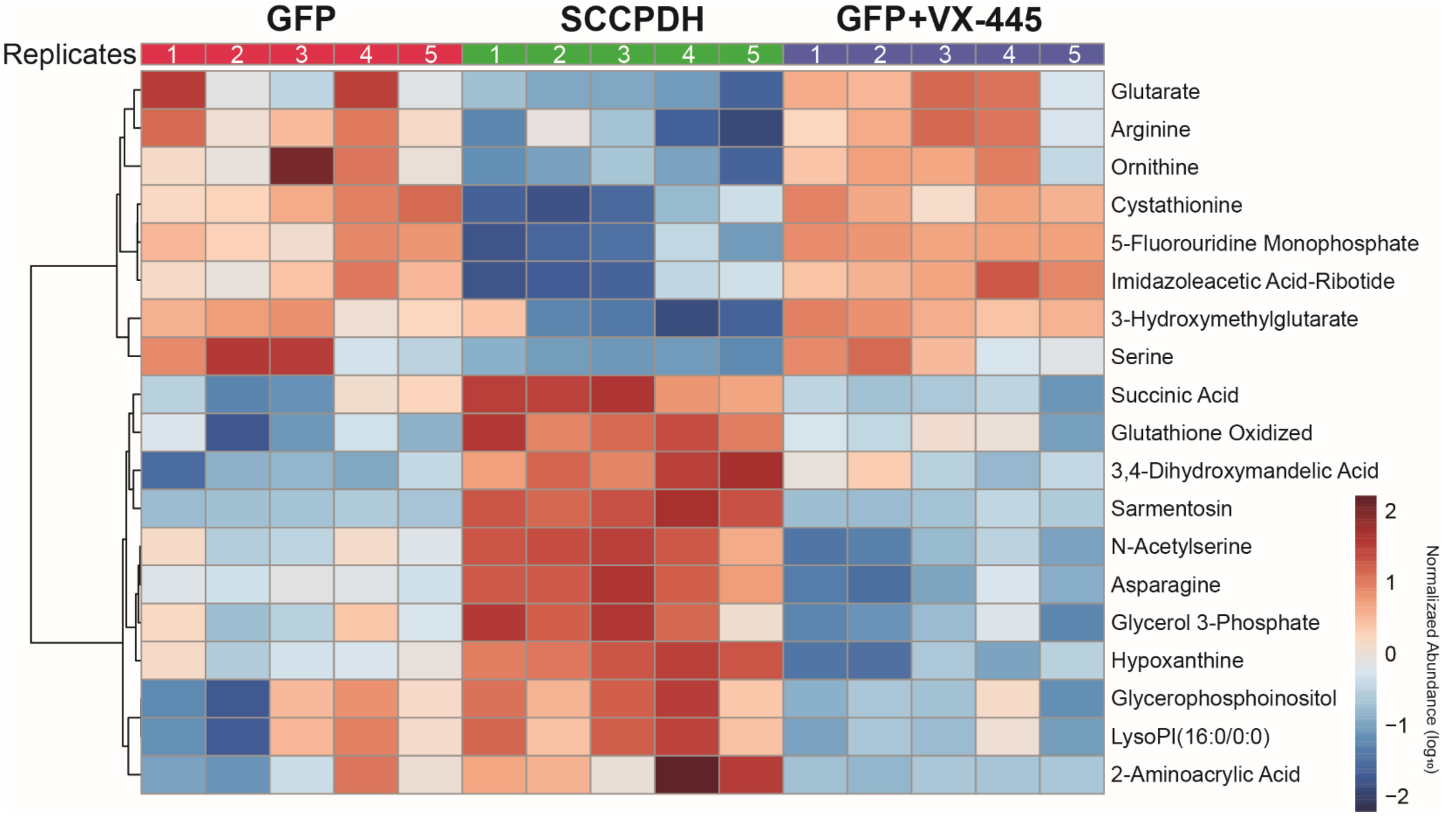


**Figure S3. Significantly changed metabolites hierarchically clustered.** Heatmap showing normalized expression of filtered metabolites within individual replicates (n = 5). Metabolites were hierarchically clustered using Pearson distance and average method.

### 1: Cysteine and methionine metabolism

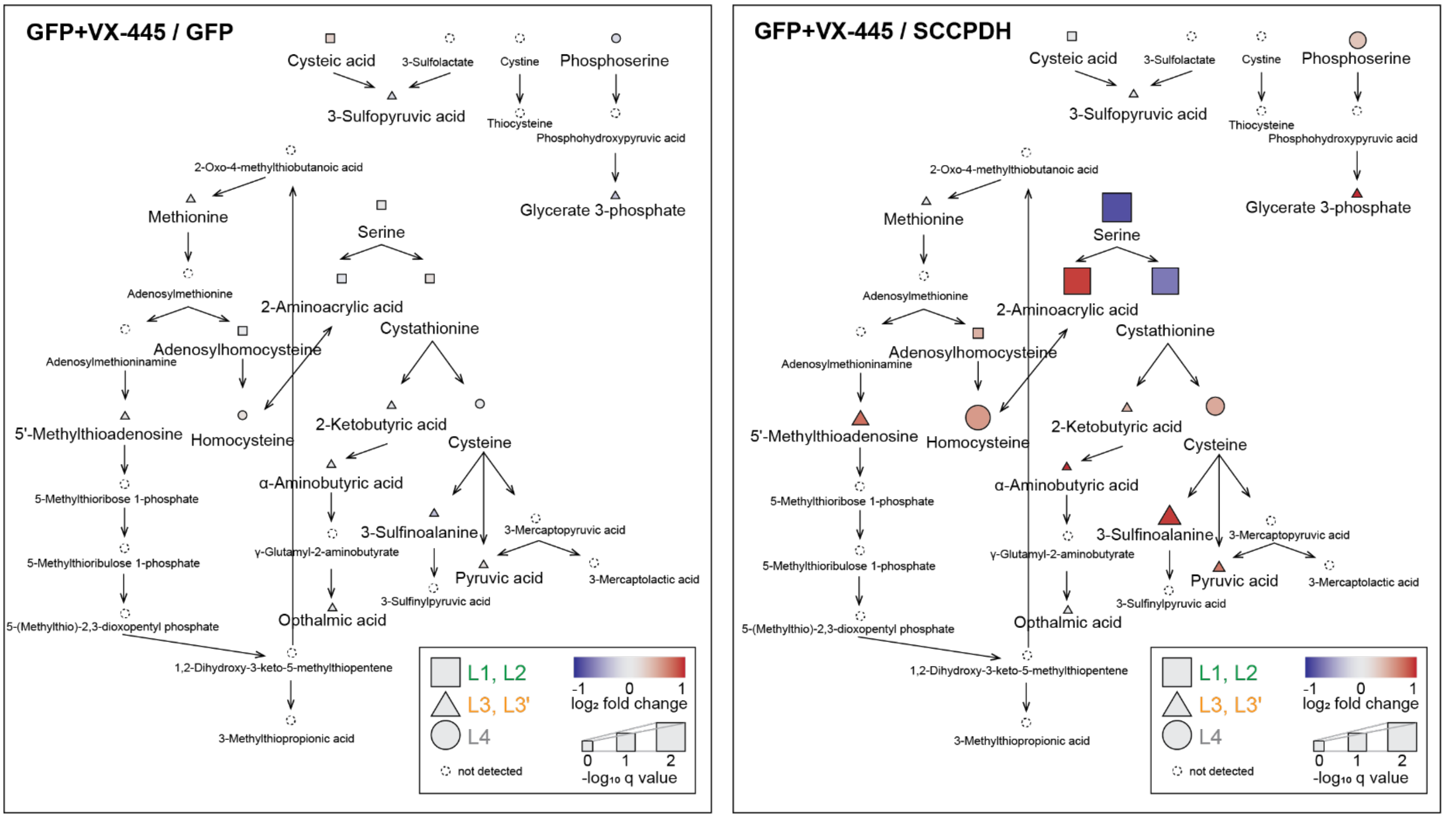


**Figure S4. Cysteine and methionine metabolism pathway (1).** Comparisons against GFP+VX-445 are shown. GFP+VX-445/SCCPDH comparison highlights the impact of VX-445 on the metabolome.

### 2: Arginine biosynthesis

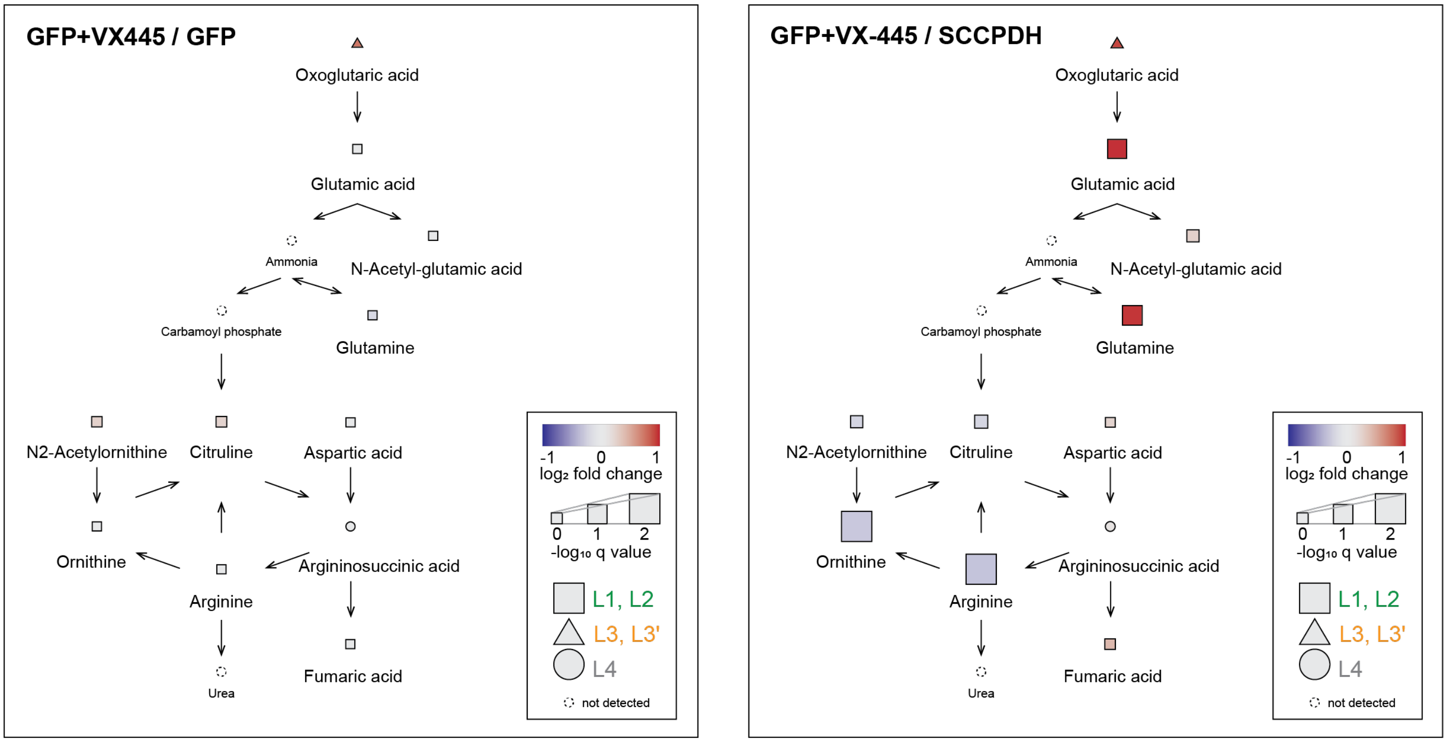


**Figure S5. Arginine biosynthesis pathway (2).** Comparisons against GFP+VX-445 are shown. GFP+VX-445/SCCPDH comparison highlights the impact of VX-445 on the metabolome.

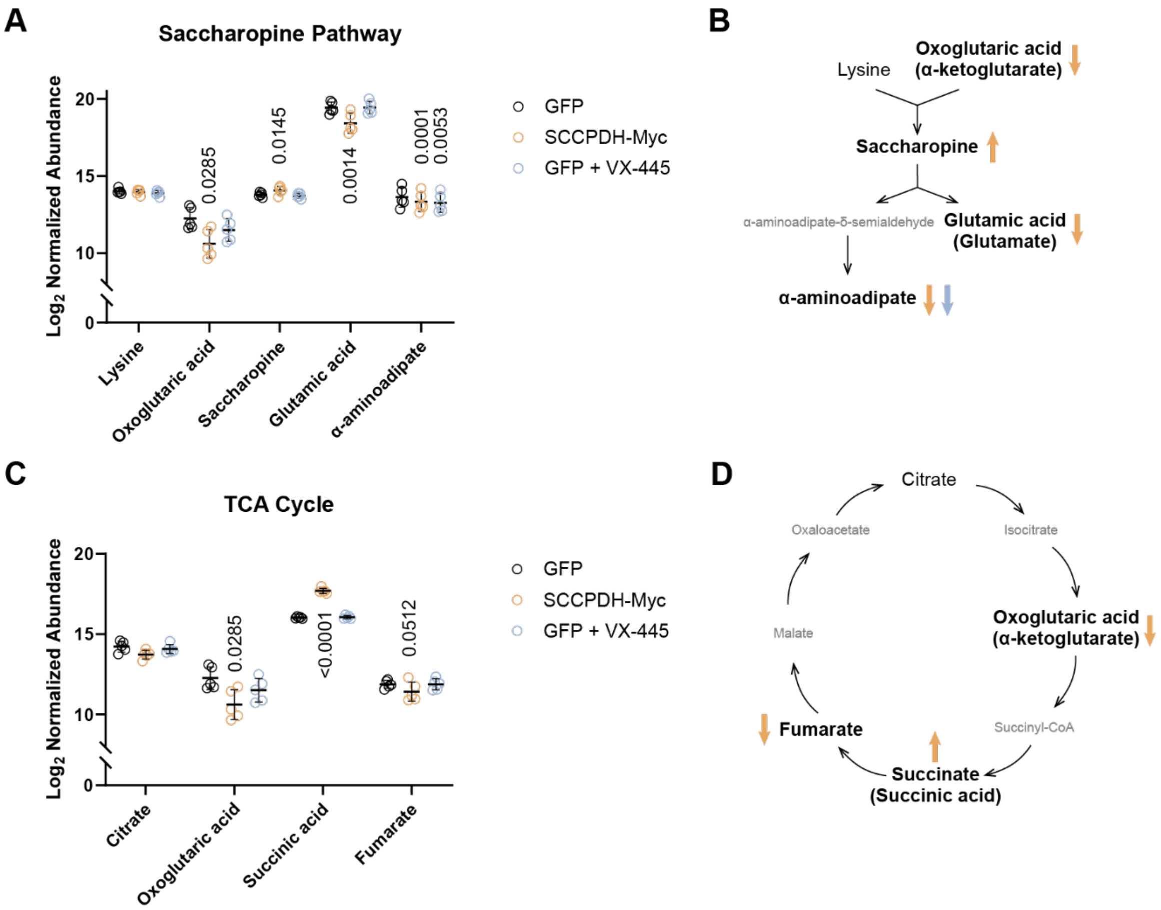


**Figure S6. Metabolites in the saccharopine pathway and TCA cycle are altered by SCCPDH overexpression.**

**A.** Metabolite abundances involved in the saccharopine pathway detected in our dataset. Statistically significant differences from paired t-tests compared to GFP are shown with p values.

**B.** Saccharopine pathway diagram. Metabolites with statistically significant changes in abundance are bolded and marked with arrows indicating direction of change (colored as in A). Metabolites not detected in our dataset are shown in gray.

**C.** Metabolite abundances involved in the TCA cycle detected in our dataset. Statistically significant differences from paired t-tests compared to GFP are shown with p values.

**D.** TCA cycle diagram. Metabolites with statistically significant changes in abundance are bolded and marked with arrows indicating direction of change (colored as in C). Metabolites not detected in our dataset are shown in gray.

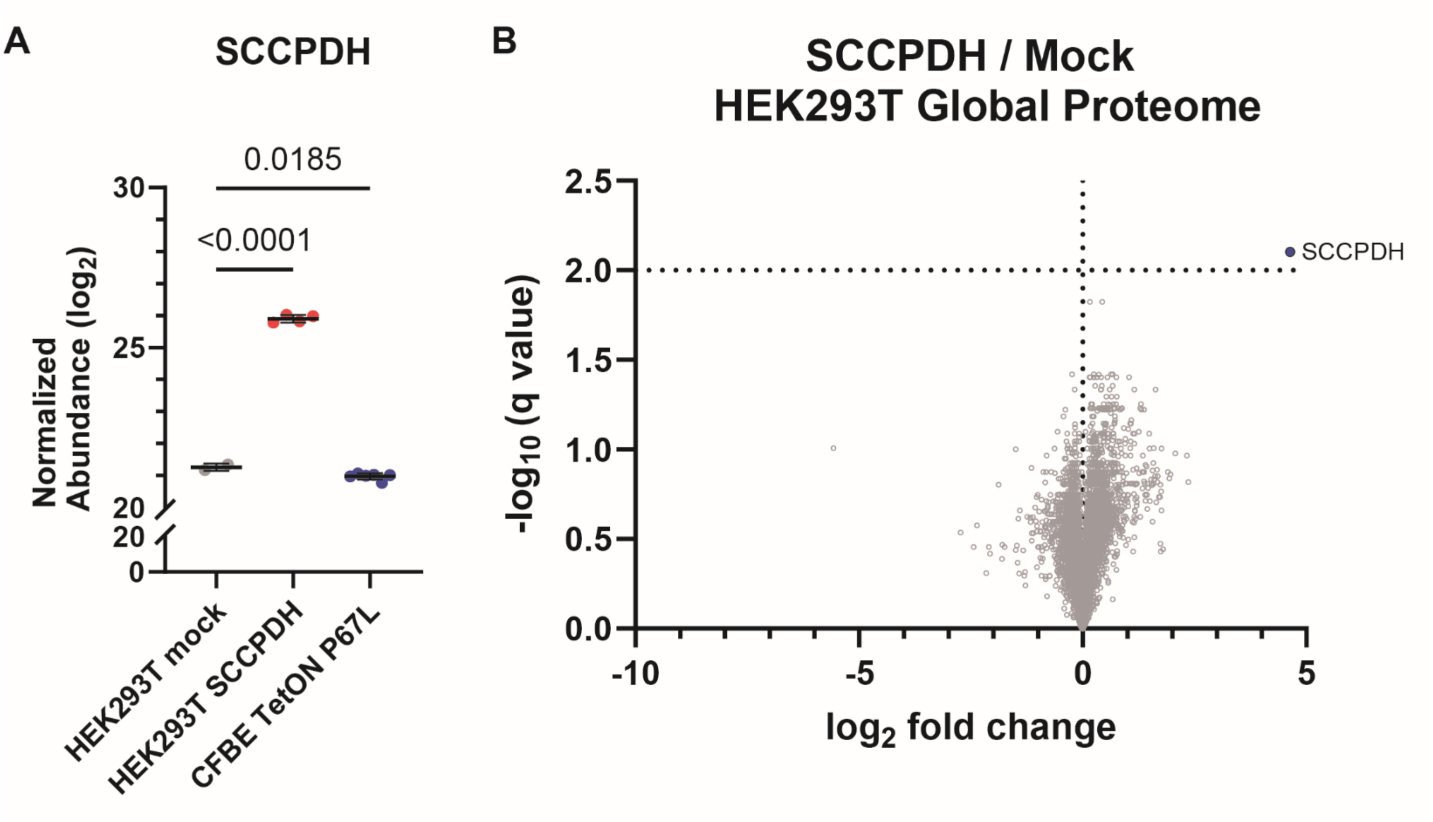


**Figure S7. Global proteomics comparison of SCCPDH expression.**

**A.** SCCPDH transfection in HEK293T cells resulted in ~32 fold change in SCCPDH abundance as determined by global proteome DIA-MS. CFBE cells showed a slight decrease in endogenous SCCPDH expression compared to HEK293T cells. Statistical differences were computed via one-way ANOVA with Geisser-Greenhouse correction and post-hoc Dunnett's multiple comparisons testing against HEK293T mock. P-values as shown ($n = 2\text{-}6$).

**B.** Comparison of global proteome between HEK293T cells transfected with SCCPDH or mock. Globally, proteins were largely unchanged. Only SCCPDH was significantly upregulated at FDR < 1% ($q < 0.01$).

**Table S3. Classification of metabolite annotations**

| ID classification | Evidence |
|---|---|
| L1 (Validated) | Precursor mass, retention time, and MS/MS data consistent with in-house library |
| L2 (Putative) | Precursor mass and MS/MS data consistent with published data |
| L3 (Tentative) | Multiple candidates possible, but a single species or group of isomers is prioritized by MS/MS data or retention time ( ' = isotopic similarity) |
| L4 (Unique Feature) | MS data |

(MS: Mass Spectrometry)

**Table S4. All metabolic pathways found from pathway analysis.**

| Metabolic Pathways | Total | Hits | -log10(p) | Impact | Components |
|---|---|---|---|---|---|
| 1 Cysteine and methionine metabolism | 33 | 3 | 2.7898 | 0.2004 | Cystathionine, Serine, Dehydroalanine |
| 2 Arginine biosynthesis | 14 | 2 | 2.3368 | 0.1371 | Arginine, Ornithine |
| 3 Alanine, aspartate and glutamate metabolism | 28 | 2 | 1.7441 | 0 | Asparagine, Succinate |
| 4 Glutathione metabolism | 28 | 2 | 1.7441 | 0.027 | Glutathione disulfide, Ornithine |
| 5 Glycine, serine and threonine metabolism | 33 | 2 | 1.6082 | 0.2146 | Serine, Cystathionine |
| 6 Arginine and proline metabolism | 36 | 2 | 1.537 | 0.2884 | Arginine, Ornithine |
| D-Amino acid metabolism | 15 | 1 | 0.96319 | 0 | Serine |
| Butanoate metabolism | 15 | 1 | 0.96319 | 0 | Succinate |
| Glycerolipid metabolism | 16 | 1 | 0.93666 | 0.0436 | Glycerol 3-Phosphate |
| Citrate cycle (TCA cycle) | 20 | 1 | 0.84576 | 0.0327 | Succinate |
| Propanoate metabolism | 22 | 1 | 0.80736 | 0 | Succinate |
| Sphingolipid metabolism | 32 | 1 | 0.65953 | 0 | Serine |
| Glyoxylate and dicarboxylate metabolism | 32 | 1 | 0.65953 | 0.0423 | Serine |
| Glycerophospholipid metabolism | 36 | 1 | 0.61431 | 0.0809 | Glycerol 3-Phosphate |
| Drug metabolism - other enzymes | 39 | 1 | 0.58398 | 0.0666 | 5-Fluorouridine monophosphate |
| Tyrosine metabolism | 42 | 1 | 0.55622 | 0.0067 | 3,4-Dihydroxymandelate |
| Purine metabolism | 70 | 1 | 0.37509 | 0.0162 | Hypoxanthine |

**Synthesis of Photoaffinity Ligand VU439**

**Materials**: Solvents were obtained from commercial or an MBraun MB-SPS solvent system. Commercial reagents were used as received.

**Instrumentation**: The preparative reverse phase HPLC was conducted on a Gilson HPLC system using a Phenomenex Luna column (100 Å, 50 x 21.20 mm, 5 µm C18) with UV/Vis detection. The compounds were purified by Teledyne ISCO normal phase column chromatograph system. $^1$H NMR spectra were recorded on Bruker 400 MHz spectrometer and are reported relative to deuterated solvent signals. Data for $^1$H NMR spectra are reported as follows: chemical shift (δ ppm), multiplicity (s = singlet, d = doublet, t = triplet, q = quartet, quint = quintet, m = multiplet, br = broad, app = apparent), coupling constants (Hz), and integration. $^{13}$C NMR spectra were recorded on Bruker 100, 125, or 150 MHz spectrometers and are reported relative to deuterated solvent signals. LC/MS was conducted and recorded on an Agilent Technologies 6130 Quadrupole instrument.

### Compound preparation:

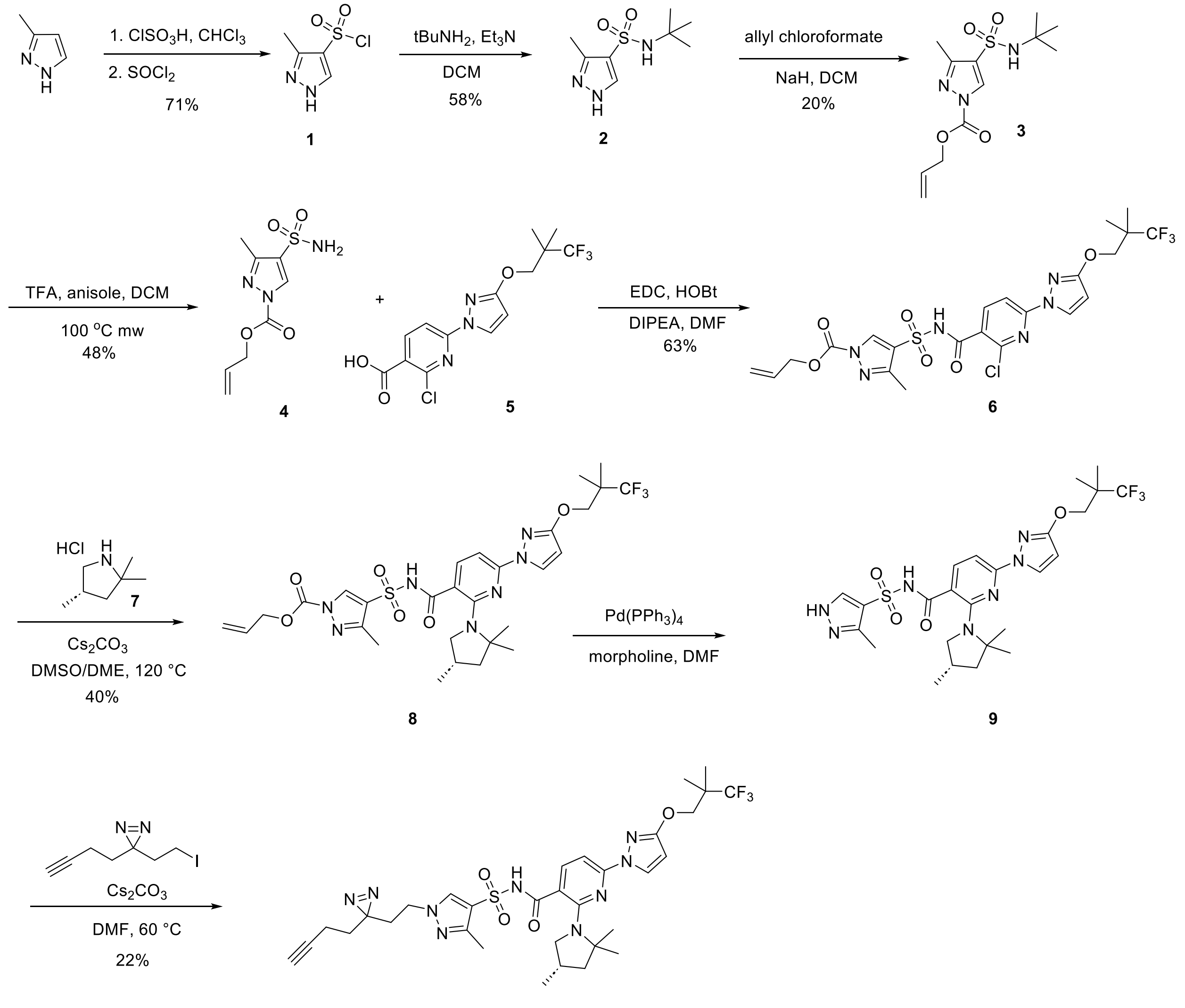


**VU0945439 (NN-445)**

**Compound 1.** To a solution of chlorosulfuric acid (23.8 mL, 355.0 mmol) in chloroform (100 mL) was added 3-methyl-1H-pyrazole (5.3 g, 64.5 mmol) in chloroform (30 mL) dropwise at 0 °C. The reaction mixture was stirred for 14 hr at 60 °C, cooled to room temperature, quenched with ice water (100 mL), and extracted with DCM (3 x 50 mL). The combined organic layers were dried over $MgSO_4$, filtered, and concentrated *in vacuo*. The crude was dissolved in hot EtOAc (50 mL) and hexanes were slowly added until white solid product was formed and washed with hexanes to afford the white solid title compound (5.36 g, 71%); $^1$H NMR (400 MHz, $CDCl_3$) δ 8.10 (s, 1H), 2.64 (s, 3H); LCMS ESI-MS(m/z) calc'd for $C_4H_5ClN_2O_2S$ $[M+H]^+$: 180.98 measured 181.0.

**Compound 2.** To a solution of **1** (4.3 g, 23.8 mmol) in DCM (80 mL) was added t-butylamine (20.1 mL, 190.5 mmol) at room temperature. The reaction mixture was stirred for 5 hr and solvent was removed *in vacuo*. The crude product was purified by ISCO column chromatography eluting with 60 to 80% EtOAc in DCM to afford the white solid title compound **2** (3.0 g, 58%); $^1$H NMR (400 MHz, $CDCl_3$) δ 7.88 (s, 1H), 2.49 (s, 3H), 1.29 (s, 9H); LCMS ESI-MS(m/z) calc'd for $C_8H_{15}N_3O_2$ $[M+H]^+$: 218.09 measured 218.1.

**Compound 3.** To a solution of **2** (1.62 g, 7.45 mmol) in DMF (25 mL) was added NaH (0.39 g, 9.7 mmol, 60% dispersion in mineral oil) at 0 °C. After stirring for 30 min at 0 °C, allyl chloroformate (0.87 mL, 8.5 mmol) was added to reaction mixture. The reaction mixture was warmed up to room temperature, stirred for 3 hr, quenched with sat. $NH_4Cl$ (50 mL), extracted with EtOAc (3 x 30 mL). The combined organic layers were dried over $MgSO_4$, filtered, and concentrated *in vacuo*. The crude product was purified by ISCO column chromatography eluting with 0 to 50% EtOAc in hexanes to afford the yellow oil compound **3** (0.45 g, 20%); $^1$H NMR (400 MHz, $CDCl_3$) δ 8.48 (s, 1H), 6.08-5.99 (m, 1H), 5.48 (dd, *J* = 17.2, 1.2 Hz, 1H), 5.40 (dd, *J* = 10.4, 0.8 Hz, 1H), 4.95 (d, *J* = 6.4 Hz, 2H), 2.48 (s, 3H), 1.29 (s, 9H).

**Compound 4.** A mixture of **3** (200 mg, 0.66 mmol), TFA (1.02 mL, 1.32 mmol), and anisole (1.5 mL, 1.32 mmol) in DCM (13.2 mL) was irradiated at 100 °C for 2 hr under microwave. The reaction mixture was concentrated in vacuo and purified by ISCO column chromatography eluting with 0 to 15% MeOH in DCM to afford the brown oil compound **4** (77 mg, 48%); $^1$H NMR (400 MHz, MeOH-$d_4$) δ 8.55 (s, 1H), 6.14-6.05 (m, 1H), 5.50 (dd, *J* = 17.2, 1.6 Hz, 1H), 5.39 (dd, *J* = 10.4, 1.2 Hz, 1H), 4.97 (d, *J* = 5.6 Hz, 2H), 2.48 (s, 3H); LCMS ESI-MS(m/z) calc'd for $C_8H_{11}N_3O_4S$ $[M+H]^+$: 246.05 measured 246.1.

**Compound 6.** A mixture of **4** (170 mg, 0.46 mmol), **5** (200 mg, 0.56 mmol, prepared the previously published procedure by Abela, A. R., et al, WO 107100, 2018), EDC (265 mg, 1.38 mmol), and HOBt (21 mg, 0.14 mmol) were suspended in DMF (4 mL) and DIPEA (0.80 mL, 4.6 mmol) was added to reaction mixture. The reaction mixture was stirred for 12 hr at room temperature, quenched with water (3 mL), and extracted with EtOAc (3 x 5 mL). The combined organic layers were dried over $MgSO_4$, filtered, and concentrated *in vacuo*. The crude product was purified by ISCO column chromatography eluting with 0 to 20% MeOH in DCM to afford the white solid title compound **6** (170 mg, 63%); $^1$H NMR (400 MHz, $CDCl_3$) δ 8.75 (s, 1H), 8.27 (d, *J* = 2.8 Hz, 1H), 8.22 (d, *J* = 8.4 Hz, 1H), 7.67 (t, *J* = 8.4 Hz, 1H), 6.00-5.92 (m, 1H), 5.97 (d, *J* = 2.8 Hz, 1H), 5.43 (d, *J* = 16.8 Hz, 1H), 5.35 (d, *J* = 10.4 Hz, 1H), 4.89 (d, *J* = 6.0 Hz, 2H), 2.99 (s, 2H), 2.54 (s, 3H), 1.26 (S, 6H); LCMS ESI-MS(m/z) calc'd for $C_{22}H_{22}ClF_3N_6O_6S$ $[M+H]^+$: 591.1 measured 591.1.

**Compound 8.** A mixture of **6** (158 mg, 0.27 mmol), (S)-2,2,4-trimethylpyrrolidine hydrochloride **7** (120 mg, 0.80 mmol), cesium carbonate (0.52 g, 1.60 mmol) in DMSO (3 mL) and DME (0.75 mL) stirred at 120 °C for 12 hr. The reaction was cooled to room temperature, diluted with DCM, acidified with 1N solution of HCl (adjusted to pH ~1), extracted with DCM (3 x 10 mL). The combined organic layers were dried over $MgSO_4$, filtered, and concentrated *in vacuo* to afford the yellow oil compound 8 (72 mg, 40%). The crude product was used in the next step without further purification.

**Compound 9.** A mixture of **8** (53 mg, 0.08 mmol), $Pd(PPh_3)_4$ (2 mg, 0.016 mmol), and morpholine (0.014 mL, 0.16 mmol) in DMF (1 mL) was sparged with Argon for 5 min, and stirred at room temperature for 5 hr. The reaction was filtered through a celite pad and washed with EtOAc. The filtrate was washed with sat. $NH_4Cl$ (3 mL), extracted with EtOAc (3 x 5 mL), and the combined organic layers were dried over $MgSO_4$, filtered, and concentrated *in vacuo.* The crude product was purified by ISCO column chromatography eluting with 0 to 40% EtOAc in DCM to afford the oil compound **9** (19 mg, 40%); $^1$H NMR (400 MHz, $CDCl_3$) δ 8.31 (d, *J* = 8.4 Hz, 1H), 8.22 (d, *J* = 2.8 Hz, 1H), 8.04 (s, 1H), 7.55 (d, *J* = 8.4 Hz, 1H), 5.97 (d, *J* = 2.8 Hz, 1H), 4.24 (s, 2H), 3.45 (t, *J* = 9.6 Hz, 1H), 3.09 (t, *J* = 8.4 Hz, 1H), 2.61 (s, 3H), 2.11 (dd, *J* = 12.4, 8.0 Hz, 1H), 1.70 (dd, *J* = 12.4, 9.6 Hz, 1H), 1.37 (s, 3H), 1.32 (s, 3H), 1.27 (s, 6H), 1.20 (d, *J* = 6.8 Hz, 3H), 1.05 (dd, *J* = 14.4, 6.4 Hz, 1H); LCMS ESI-MS(m/z) calc'd for $C_{25}H_{32}F_3N_7O_4S$ $[M+H]^+$: 584.22 measured 584.3.

**VU0945439 (VU439).** To a solution of **9** (19 mg, 0.033 mmol) in DMF (0.5 mL) was added cesium carbonate (33 mg, 0.1 mmol), followed by 3-(but-3-yn-1-yl)-3-(2-iodoethyl)-3H-diazirine (16 mg, 0.065 mmol). The reaction was stirred for 12 hr at 50 °C, quenched with 1N solution of citric acid (3 mL), diluted with EtOAc (10 mL), washed with water (5 mL) and brine (3 mL). The combined organic layers were dried over $MgSO_4$, filtered, and concentrated *in vacuo*. The crude product was purified by ISCO column chromatography eluting with 0 to 30% EtOAc in DCM to afford the title compound **VU0945439 (VU439,** 5mg, 22%);$^1$H NMR (400 MHz, $CDCl_3$) δ 8.34 (dd, *J* = 8.4, 1.2 Hz, 1H), 8.23 (d, *J* = 2.8 Hz, 1H), 8.15 (s, 1H), 7.59 (d, *J* = 8.8 Hz, 1H), 5.98 (d, *J* = 2.8 Hz, 1H), 4.24 (s, 2H), 3.92 (t, *J* = 7.2 Hz, 2H), 3.52 (t, *J* = 9.2 Hz, 1H), 3.14-3.05 (m, 1H), 2.71 (s, 1H), 2.69-2.60 (m, 1H), 2.48 (s, 3H), 2.14 (dd, *J* = 12.4, 8.0 Hz, 1H), 1.99-1.92 (m, 4H), 1.71 (m, 1H), 1.56-1.50 (m, 2H), 1.36 (s, 3H), 1.31 (s, 3H), 1.27 (s, 6H), 1.21 (d, *J* = 6.8 Hz, 3H); LCMS ESI-MS(m/z) calc'd for $C_{32}H_{40}F_3N_9O_4S$ $[M+H]^+$: 704.3 measured 704.7.

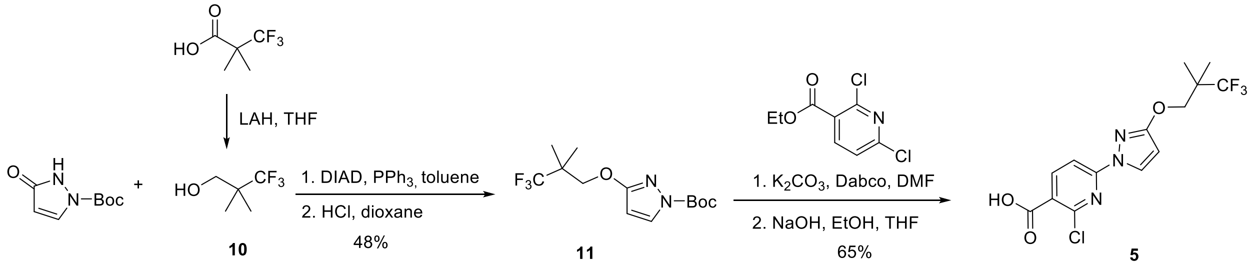


**Compound 10**. To a solution of 3,3,3-trifluoro-2,2-dimethylpropanoic acid (2.0g, 12.8 mmol) in $Et_2O$ (50 mL) was cooled to 0°C. $LiAlH_4$ (0.63g, 16.7mmol) was added portion wise over 15 min. This was then allowed to stir at 0°C, for 1.5 hr, then to warm to room temperature slowly and stir for 16 hr. At this time the reaction was again cooled to 0°C, where $H_2O$ (1 mL) was added followed by NaOH (2M, 1 mL) and finally $H_2O$ (3 mL). This was then allowed to stir at 0°C for 30 min. The reaction was then filtered over a celite pad. This was then concentrated under gentle vacuum to provide a clear colorless oil (1.86 g) containing a mixture of the product

compound **10** in $Et_2O$ (63 % weight of product ~1.17g, and 37 wt.% $Et_2O$ as determined by 1H-NMR). $^1H$ NMR (400 MHz, $CDCl_3$) δ 3.62 (d, *J* = 6.4 Hz, 2H), 1.16 (s, 6H);

***tert*-Butyl 3-(3,3,3-trifluoro-2,2-dimethylpropoxy)-1*H*-pyrazole-1-carboxylate.** A mixture of 3,3,3-trifluoro-2,2-dimethylpropan-1-ol (1.5g, 10.6 mmol) and 1-Boc-1*H*-pyrazol-3(2*H*)-one (1.94g, 10.6 mmol) in toluene (21 mL) was added triphenyl phosphine (3.04g, 11.6 mmol) followed by DIAD (2.28 mL, 11.6). The resulting mixture was then allowed to stir at 110°C for 16 hr. The mixture was then allowed to cool to R.T. and concentrated *in vacuo* to an oil. To the crude material was added heptane (30 mL) This was allowed to stir vigorously for 2 hr. The solid triphenyl phosphine oxide was removed by filtration and washed with a mixture of heptane: toluene (4:1, 100 mL). The crude material was then concentrated *in vacuo.* The crude product was purified by ISCO column chromatography eluting with 0 to 40% EtOAc in Hexane to afford tert-butyl 3-(3,3,3-trifluoro-2,2-dimethylpropoxy)-1H-pyrazole-1-carboxylate (1.62g, 49%) $^1H$ NMR (400 MHz, $CDCl_3$) δ 7.86 (d, *J* = 2.9 Hz, 1H), 5.92 (d, *J* = 2.9 Hz, 1H), 4.28 (s, 2H), 1.64 (s, 9H), 1.26 (s, 6H); LCMS ESI-MS(m/z) calc'd for $C_9H_{11}F_3N_2O_2$ $[M\text{-}tBu +H]^+$: 253.07 measured 253.1.

**Compound 11**. To a solution of *tert*-butyl 3-(3,3,3-trifluoro-2,2-dimethylpropoxy)-1*H*-pyrazole-1-carboxylate (1.6g, 5.2 mmol) was added HCl in dioxane (4M, 6.5mL, 26.0mmol). This was then heated at 45°C for 2h. This was then allowed to cool to R.T. and concentrated *in vacuo* to an oil. This was then diluted with NaOH (1M aq, 10 mL) and extracted with $Et_2O$ (3 x 15 mL). The combined organics were washed with brine and dried over $MgSO_4$, filtered, and concentrated *in vacuo* to give product (1.05g, 97%). $^1H$ NMR (400 MHz, $CDCl_3$) δ 7.39 (d, *J* = 2.5 Hz, 1H), 5.78 (d, J = 2.5 Hz, 1H), 4.15 (s, 2H), 3.73 (s, 1H), 1.28 (s, 6H); LCMS ESI-MS(m/z) calc'd for $C_8H_{11}F_3N_2O$ $[M+H]^+$: 208.08 measured 209.1.

**Ethyl 2-chloro-6-(3-(3,3,3-trifluoro-2,2-dimethylpropoxy)-1*H*-pyrazol-1-yl)nicotinate.** To a solution of ethyl 2,6-dichloronicotinate (1.2g, 5.1 mmol) in DMF (12.7 mL) was added 3-(3,3,3-trifluoro-2,2-dimethylpropoxy)-1*H*-pyrazole (1.06g, 5.1mmol), $K_2CO_3$ (910 mg, 6.6mmol) and DABCO (86 mg, 0.76 mmol). This was then allowed to stir at R.T. for 16hr. The reaction was then cooled to 0°C and allowed to stir for 10 min. At this time $H_2O$ (20mL) was added and allowed to stir at 0°C for 2 hr. The reaction was filtered to remove the solid and washed with $H_2O$ (400mL) to afford the title compound (1.8g, 90%). $^1H$ NMR (400 MHz, $CDCl_3$) δ 8.40 (d, *J* = 2.8 Hz, 1H), 8.29 (d, *J* = 8.5 Hz, 1H), 7.74 (d, *J* = 8.5 Hz, 1H), 6.02 (d, *J* = 2.8 Hz, 1H), 4.43 (q, *J* = 7.1 Hz, 2H), 4.27 (s, 2H), 1.44 (t, *J* = 7.08 Hz, 3H), 1.30 (s, 6H). LCMS ESI-MS(m/z) calc'd for $C_{16}H_{18}ClF_3N_3O_3$ $[M+H]^+$: 393.09 measured 392.1.

**Compound 5.** To a solution of ethyl 2-chloro-6-(3-(3,3,3-trifluoro-2,2-dimethylpropoxy)-1*H*-pyrazol-1-yl)nicotinate (1.8g, 4.59mmol) in MeOH:THF (1:1, 100mL) was added NaOH (2M aq, 4.6 mL). This was allowed to stir at R.T. for 2hr. At this time the reaction was diluted to HCl (1M aq, 5mL), and $H_2O$ (2.5 mL).

The reaction was then extracted with EtOAc (3 x 50mL). The combined organics were dried over $MgSO_4$, filtered, and concentrated *in vacuo* to give the title compound as a solid (1.2g, 72%). $^1$H NMR (400 MHz, DMSO-d$_6$) δ 8.46 (d, $J$ = 2.8 Hz, 1H), 8.40 (d, $J$ = 8.5 Hz, 1H), 7.77 (d, $J$ = 8.5 Hz, 1H), 6.26 (d, $J$ = 2.8 Hz, 1H), 4.28 (s, 2H), 1.24 (s, 6H).

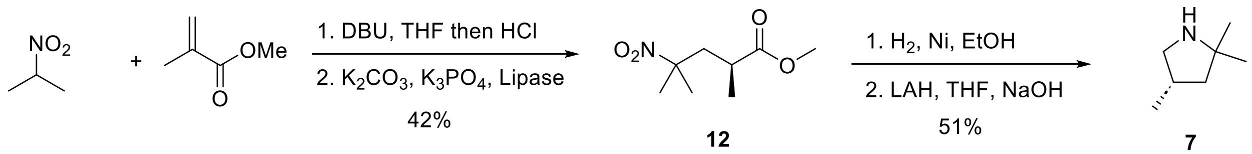

12 7

**Methyl 2,4-dimethyl-4-nitropentanoate**. DBU (4.17 mL, 27.9 mmol) was added to a solution of nitropropane (5 mL, 55.7 mmol) in THF (25 mL) and stirred at 50 °C. Methyl methacrylate (6.53 mL, 69.4 mmol) in THF (10 mL) was added to a dropping funnel and slowly added over 30 min. The reaction mixture was stirred at 50 °C for 20 hr. The reaction mixture was concentrated, and the residue was dissolved in $Et_2O$ (60 mL). 2N HCl solution was added (adjusted to pH 2). Extracted with $Et_2O$ (4 x 25 mL) and the combined organic layers were dried over $MgSO_4$, filtered, and concentrated *in vacuo.* The crude mixture (8.89 g, 85%) was used in the next step without further purification; $^1$H NMR (400 MHz, $CDCl_3$) δ 3.67 (s, 3H), 2.52-2.42 (m, 2H), 2.04 (dd, $J$ = 13.8, 1.8 Hz, 1H), 1.58 (s, 3H), 1.53 (s, 3H), 1.19 (d, $J$ = 6.9 Hz, 3H).

**Compound 12.** Monopotassium phosphate (1.15 g, 8.45 mmol) and lipase (3.80 g) were added to a solution of methyl 2,4-dimethyl-4-nitropentanoate (8.89 g, 46.9 mmol) in $H_2O$ (50 mL). 20% aq. $K_2CO_3$ was added to adjust pH ~6.5. The reaction mixture was stirred at 32 °C for 21 hr. The reaction mixture was extracted with $Et_2O$ (3 x 50 mL) and the combined organic layers were washed with 10% aq. $Na_2CO_3$ (x 3) and brine. The organic layer was filtered through a Celite pad (washed with EtOAc) and the liquid was extracted with (3 x 20 mL). The combined organic layers were dried over $MgSO_4$, filtered, and concentrated *in vacuo.* The crude mixture (4.37 g, 49%) was used in the next step without further purification; $^1$H NMR (400 MHz, $CDCl_3$) δ 3.67 (s, 3H), 2.54-2.40 (m, 2H), 2.04 (dd, $J$ = 13.8, 2.3 Hz, 1H), 1.58 (s, 3H), 1.53 (s, 3H), 1.18 (d, $J$ = 6.8 Hz, 3H).

**(*S*)-3,5,5-Trimethylpyrrolidin-2-one**. Raney-Ni (1.50 g) was added to a solution of methyl (*S*)-2,4-dimethyl-4-nitropentanoate (4.37 g, 23.1 mmol) in ethanol (150 mL). The reaction mixture was degassed with Ar(g) for 15 min and stirred at 60 °C with $H_2$(g) for 4 hr and at 50 °C for 18 hr. The reaction mixture was filtered through a Celite/$MgSO_4$ pad (washed with ethanol) and concentrated *in vacuo.* The crude mixture (2.84 g, 97%) was used in the next step without further purification; $^1$H NMR (400 MHz, $CDCl_3$) δ 5.70 (br s, 1H), 2.66-2.57 (m, 1H), 2.17 (dd, $J$ = 12.6, 8.8 Hz, 1H), 1.56 (dd, $J$ = 12.6, 10.0 Hz, 1H), 1.30 (s, 3H), 1.25 (s, 3H), 1.20 (d, $J$ = 7.2 Hz, 3H); LCMS ESI-MS(m/z) calc'd for $C_7H_{13}NO$ $[M+H]^+$: 128.1 measured 128.1.

**Compound 7**. Lithium aluminum hydride (1.02 g, 26.9 mmol) was added to a solution of (*S*)-3,5,5-trimethylpyrrolidin-2-one (2.84 g, 22.3 mmol) in THF (100 mL) at 0 °C. The reaction mixture was stirred at 60 °C for 20 hr. Additional lithium aluminum hydride (800 mg, 21.1 mmol) was added at 0 °C. The reaction mixture was stirred at 60 °C for 3 hr. Additional lithium aluminum hydride (800 mg, 21.1 mmol) was added at 0 °C. The reaction mixture was stirred at 65 °C for 18 hr. The reaction was quenched with $H_2O$ (4 mL) and 50% aq. NaOH solution (15 mL) at 0 °C. The mixture was stirred at 0 °C for 1 hr and filtered through a Celite pad (washed with THF). The filtrate was dried over $MgSO_4$ and filtered. The filtrate was cooled at 0 °C and Conc. HCl (3 mL, 37% w/w) was added slowly. The mixture was stirred for 30 min at 0 °C and concentrated *in vacuo*. Toluene (4 mL) was added and concentrated *in vacuo* (repeat 3 times). Isopropanol (6 mL) was added and kept at 50 °C for 5 min. $Et_2O$ (20 mL) was added and cooled to 0 °C for 20 min. The desired product was collected by filtration (1.78 g, 53%); $^1$H NMR (400 MHz, DMSO-$d_6$) δ 9.04 (br s, 1H), 3.38-3.30 (m, 1H), 2.78-2.73 (m, 1H), 2.49-2.43 (m. 1H), 1.98 (dd, *J* = 13.0, 7.9 Hz, 1H), 3.18 (s, 3H), 1.37-1.32 (m, 1H), 2.94 (s, 3H), 1.04 (d, *J* = 6.7 Hz, 3H); LCMS ESI-MS(m/z) calc'd for $C_7H_{15}N$ $[M+H]^+$: 114.1 measured 114.2.
